# Functional Organization of the Neonatal Basal Ganglia and Thalamus

**DOI:** 10.64898/2026.07.02.736181

**Authors:** Samantha L. Blake, Jeanette K. Kenley, Tara A. Smyser, Aidan Latham, Dimitrios Alexopoulos, Deanna J. Greene, Rachel E. Lean, Deanna M. Barch, Barbara B. Warner, Joan L. Luby, Cynthia E. Rogers, Christopher D. Smyser, Chad M. Sylvester, Ashley N. Nielsen

## Abstract

The basal ganglia and thalamus are key nodes in subcortico-cortical loops involved in sensory, motor, and cognitive function. In adults, posterior regions of the subcortex link to cortical sensorimotor networks and anterior regions link to association networks. Alterations in the size, strength, and selectivity of these subcortical regional network representations are implicated in several neuropsychiatric disorders, many of which originate early in development. However, the organization of these network representations at birth remains incompletely understood, limiting our ability to devise normative and atypical developmental models of subcortico-cortical interactions. Using resting-state fMRI, we characterized the size, strength, and selectivity of cortical network representations in the basal ganglia and thalamus in a set of neonates (n=261) and compared results to children (age range 9-11 years, n=69) and adults (n=120). We found that the broad anterior-posterior organization of the subcortex is present at birth, yet representations of somatomotor networks were larger at birth compared to children and adults (p<0.001). The strength and selectivity of subcortico-cortical functional connectivity (FC) exhibited interactions between age group and network (all p<0.001), such that subcortical representations of sensorimotor networks exhibited stronger FC and higher selectivity in neonates, while subcortical representations of association networks exhibited stronger FC and higher selectivity in older cohorts. In parallel, data-driven clustering revealed areas with integration of multiple networks in the neonatal subcortex. These results suggest that subcortico-cortical FC evolves over development largely in a sensorimotor-association manner and provide a baseline for normative and disordered subcortical development.

**HIGHLIGHTS:**

- The basic anterior-posterior layout of association to sensorimotor network representation in the basal ganglia and thalamus is present at term birth.
- Sensorimotor systems are overrepresented in neonates compared to children and adults.
- Sensorimotor representations exhibit greater functional connectivity strength and selectivity than association representations at birth, while association representations are stronger and more selective than sensorimotor representations in adults.
- The neonatal subcortex exhibits substantial integration of multiple cortical networks.

**GRAPHICAL ABSTRACT:** 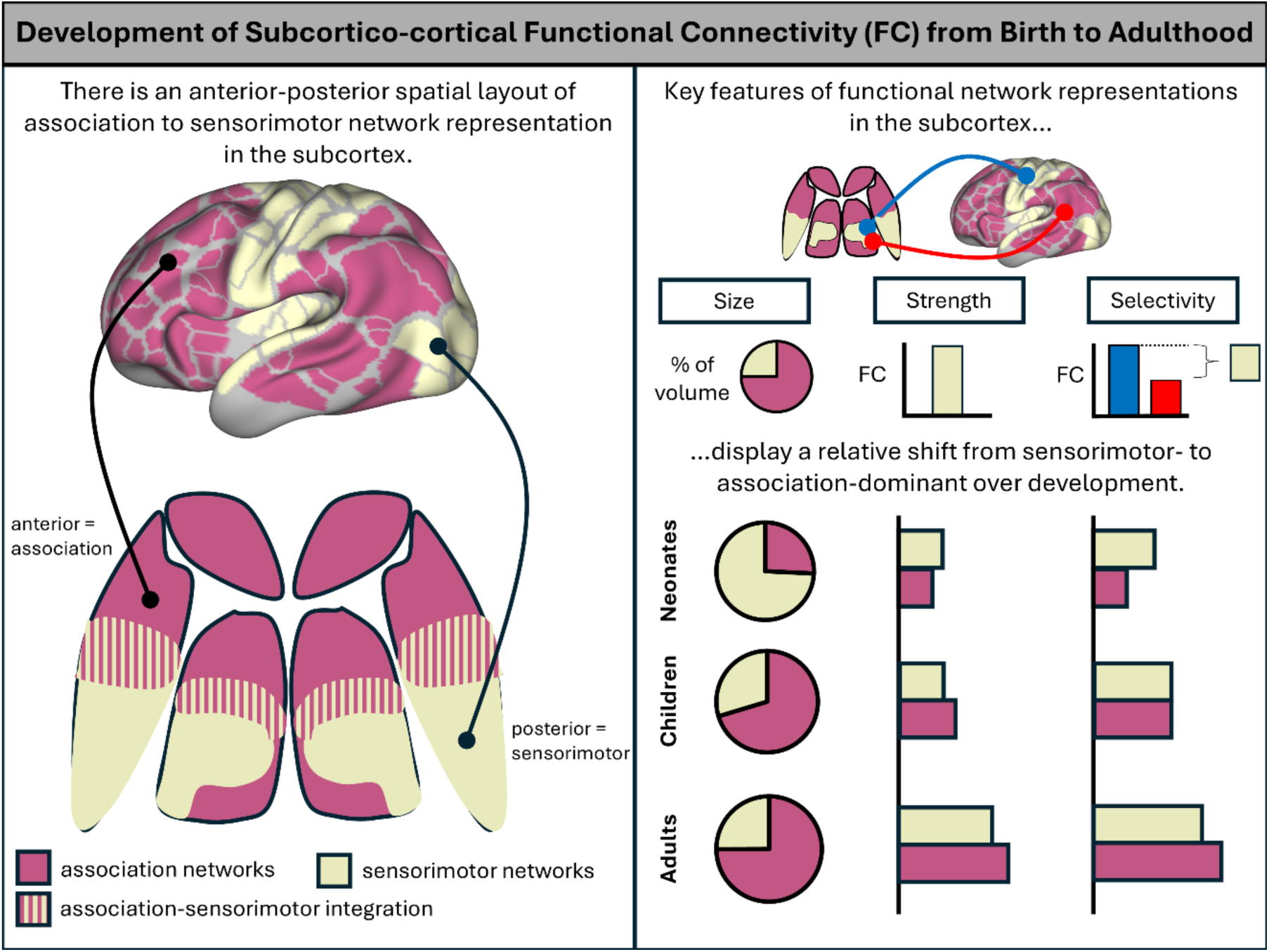

## INTRODUCTION

Neural circuits involving the basal ganglia and thalamus are central to both motor and higher cognitive function, and alterations in these subcortical structures have been linked to a host of neurological and psychiatric disorders (Nair et al., 2013; Brown et al., 2017; Ramkiran et al., 2019; Karavallil Achuthan et al., 2023). A key organizational feature is selective connectivity of specific subdivisions of the basal ganglia and the thalamus with specific portions of the cortex. In addition to this specificity, functional integration zones within the basal ganglia and thalamus connect more broadly to different parts of cortex (Haber et al., 2006; Haber and Calzavara, 2009; Haber, 2016), potentially facilitating the transfer of information between functional networks (Hwang et al., 2017).

Neuroanatomically, major thalamic nuclei and divisions in the basal ganglia can be observed prenatally (Rakić and Sidman, 1969; Kostović, 1986; Puelles and Rubenstein, 2003; Vue et al., 2009). Beginning prenatally and continuing after birth, subcortical afferents drive the functional organization of the basal ganglia and thalamus, which in turn drive the organization of their target cortical layers (LeVay and Gilbert, 1976; Govaert et al., 2025). However, the state of the functional organization of the subcortex beginning at birth is incompletely understood, limiting our ability to describe developmental cascades of sensory, motor, and cognitive function and how dysfunction in emerging subcortical functional organization may contribute to developmental disorders.

Functional connectivity (FC), the correlation in the blood oxygen level-dependent (BOLD) signal between two brain regions as measured with fMRI (Fox et al., 2005), has been used to characterize the functional organization of the basal ganglia and thalamus in humans. In children and adults, FC between the cortex and the basal ganglia (Di Martino et al., 2008; Barnes et al., 2010; Choi et al., 2012; Greene et al., 2014) and the thalamus (Fair et al., 2010; Zhang et al., 2010; Steiner et al., 2020; Huang et al., 2021; Badke D’Andrea et al., 2023) largely align with neuroanatomically-defined organization. Posterior portions of the subcortex exhibit the greatest FC to sensorimotor networks (e.g., somatomotor, auditory, visual), whereas more anterior areas exhibit greater FC to association networks (e.g., salience, cingulo-opercular, default mode, frontoparietal). Some regions within the basal ganglia and thalamus exhibit strong FC with multiple functional networks in the cortex, consistent with subcortical integration zones (Hwang et al., 2017; Greene et al., 2020; Badke D’Andrea et al., 2023). FC in children indicates that these organizing principles of subcortical functional organization are largely established by childhood (Greene et al., 2014; Badke D’Andrea et al., 2023), but whether and to what extent this organization is present at birth remains unknown.

One hypothesis is that the functional organization of the neonatal subcortex may reflect the developmental processes occurring in the cortex and the relative maturity of different neonatal cortical functional networks. During the perinatal period, several neurobiological developmental processes (e.g., axonal guidance, synaptogenesis, synaptic pruning, maturation of inhibition) occur earlier in sensorimotor systems than in association systems (Flechsig Of Leipsic, 1901; Huttenlocher and Dabholkar, 1997; Travis et al., 2005; Knowles et al., 2021; Sydnor et al., 2021). This neurodevelopmental pattern is reflected in neuroimaging findings (Sydnor et al., 2023) that characterize developmental changes in the structure and function of the sensorimotor and association cortices. The cortex expands nonlinearly such that sensorimotor cortices expand and fold earlier than association cortices, and a higher relative percentage of cortical surface area is devoted to sensorimotor cortices at birth than when fully mature (Garcia et al., 2018; Gorham et al., 2024). The strength of FC among regions within cortical functional networks, which may reflect activity-dependent Hebbian plasticity and synaptic strength (Supekar et al., 2009; Uddin et al., 2024), increases over the course of development (Thomason et al., 2013; Sylvester et al., 2023). Notably, FC strength increases at different rates such that strong within-network FC is present at birth in sensorimotor networks and not in association networks. Similarly, the selectivity of cortical FC (i.e., the difference between FC within a network and FC across networks) increases as functional networks mature and become more specialized (Nielsen et al., 2023; Sylvester et al., 2023; Tooley et al., 2024). The refinement of FC occurs at different rates and varies across the cortex, with sensorimotor networks exhibiting greater network selectivity than association networks at birth. Generally, greater network selectivity suggests decreased functional integration, which is indicated by greater between-network connectivity (Gratton et al., 2018).

Prior work investigating subcortico-cortical FC in infants and older children provides evidence in support of the hypothesis that subcortical development parallels the pattern of neurodevelopment in the cortex. Sensorimotor network representation is larger than the association network representation in the infant thalamus (Toulmin et al., 2015), and the relative representation of the sensorimotor versus salience networks decreases in the first years of life (Alcauter et al., 2014). However, less is known about the organization of the neonatal basal ganglia, the representations of other association networks in the thalamus, or how these representations change over development. From childhood to adulthood, the relative size and FC strength of sensorimotor network representations in the subcortex decrease from ages 7-21 years (Greene et al., 2014; Huang et al., 2021; Badke D’Andrea et al., 2023), while the relative size and strength of association network representations in the subcortex increase with age over that same time period (Fair et al., 2010; Huang et al., 2021). Whether these developmental trends of increased sensorimotor FC strength and selectivity predict the functional organization in the neonatal subcortex remains to be determined.

The goal of this study was to characterize the functional organization of the basal ganglia and thalamus in neonates (mean PMA 41.3 weeks) using resting-state fMRI and to determine how the size, strength, and selectivity of functional network representations in the subcortex compared to those in children (mean 9.9 years) and adults (mean 25 years). We subdivided the subcortex into regions based on the single cortical network with which they had the strongest FC. Based on the model that the subcortex would reflect the relative maturity of cortical functional networks, we hypothesized that neonatal subcortical representations of sensorimotor networks would be relatively larger and exhibit stronger and more selective FC than association networks when compared to older ages. In parallel data-driven analyses, we clustered the subcortex based on the overall pattern of FC to all cortical networks, which permitted us to characterize a set of regions containing representations of multiple functional networks and quantify the extent of integration in the neonatal subcortex. This work serves as a starting point for characterizing trajectories of both normative and atypical subcortical development beginning at birth.

## METHODS

### Participants

This work combines data from neonates, children, and adults that participated in separate studies that were approved by the Human Research Protection Office at Washington University in St. Louis. Informed consent was obtained from parents/guardians of neonatal and child participants and from adult participants. The neonatal dataset in this work was acquired through the Early Life Adversity and Biological Embedding (eLABE) study and has been previously described (Lean et al., 2022; Nielsen et al., 2023; Sylvester et al., 2023; Myers et al., 2024). The present study focused on resting-state fMRI data acquired during natural sleep from 261 healthy, full-term neonates (average postmenstrual age (PMA) at scan 41.3 wks, range 38-45; 141 males). Only data from participants with at least 10 min of low motion, whole-brain resting-state data after motion scrubbing (average 16.6 min; see below) were included in these analyses.

The child sample, WUNDER, consisted of 69 full-term born children between the ages of 9-11 years (average age: 9.9 years; 30 males). Only data from child participants with at least 5 min of resting-state fMRI data after motion scrubbing (average 15.2 min; see below) were included in these analyses. The adult sample consisted of 120 adults (19-31 years; 60 males) that were collected at Washington University (Power et al., 2011; Greene et al., 2014; Gordon et al., 2016). Only data from adult participants with at least 5 min of resting-state fMRI data after motion scrubbing (average 11.6 min; see below) were included in these analyses. Inclusion and exclusion criteria for each dataset are summarized in the **Supplementary Methods**.

### Image Acquisition

#### Neonates

Neuroimaging was performed in full-term neonates during natural sleep on a Siemens PRISMA 3T MRI scanner with a 64-channel head coil (**Supp. Methods**). A T2-weighted image (sagittal, 208 slices, 0.8-mm isotropic resolution, time to echo (TE)=563 ms, tissue T2=160 ms, repetition time (TR)=3200 ms) was collected. Functional imaging (fMRI) was performed using a BOLD-gradient-recalled echo-planar multiband (MB) sequence (72 slices, 2.0-mm isotropic resolution, TE=37 ms, TR=800 ms, MB factor=8). Using the same parameters, spin-echo field maps were obtained (at least 1 anterior-posterior and 1 posterior-anterior; **Supp. Methods**). Runs were 420 frames and ∼5.6 min in length. Framewise integrated real-time MRI monitoring (FIRMM) was used during scanning to monitor real-time neonatal movement (Dosenbach et al., 2017; Badke D’Andrea et al., 2022).

### Image Acquisition: Children and Adults

Methods for image acquisition and neuroimaging parameters for the child and adult datasets are detailed in the **Supplementary Methods**.

### Image Analysis

#### Preprocessing

For each dataset, fMRI preprocessing was conducted with minor variations to optimize data quality metrics per age group. fMRI preprocessing steps included correction of intensity differences, bias field correction, intensity normalization of each run to a whole-brain mode value of 1000, and linear registration of BOLD images to the adult Talairach isotropic atlas (Talairach and Tournoux, 1988). Volumetric data were sampled to a 3 × 3 × 3 mm space in the same step. Prior to FC processing, cortical volumetric preprocessed BOLD data were mapped to subject-specific surfaces using established procedures adapted from the Human Connectome Project (Marcus et al., 2011, 2013). See **Supplementary Methods** for further dataset-specific preprocessing steps.

#### FC Processing

Following initial preprocessing, resting-state fMRI BOLD timeseries underwent several additional preprocessing steps (Power et al., 2014; Gordon et al., 2016; Feczko et al., 2021), as detailed in the **Supplementary Methods**, including demeaning and detrending within run, multiple regression with nuisances timeseries, and bandpass filtering.

#### Motion Scrubbing

fMRI data were censored at framewise displacement (FD)>0.25mm for neonates, FD>0.3mm for children, and FD>0.2mm for adults. Because head motion and FD distributions vary across datasets based on numerous factors including head size, breathing rate, and sampling frequency (TR), we selected different FD thresholds for each dataset based on visual inspection of the FD traces (Power et al., 2014; Power, 2017). To be included in the study, neonates required a minimum of 10 min of data, and children and adults required a minimum of 5 min. Additional motion censoring requirements are detailed in the **Supplementary Methods**.

### Anatomical Definitions

#### Subcortical Definitions

Basal ganglia and thalamus segmentations for all datasets were derived from atlas-aligned data and then applied to individuals (see **Supp. Methods**). The basal ganglia mask included the caudate, putamen, and pallidum.

#### Cortical Functional Network Definition

To investigate FC between the subcortex and the cortex, we selected a subset of the most commonly identified functional networks in adults (Gordon et al., 2016). This process resulted in a set of 10 networks: (1) sensorimotor systems: visual, auditory, somatomotor face, and somatomotor hand; (2) association systems: default mode, frontoparietal, cingulo-opercular, dorsal attention, ventral attention, and salience. To ensure that our findings were not unique to these network definitions, we also tested network configurations that were defined in neonates (Myers et al., 2024) and that included more detailed somatomotor organization (Gordon et al., 2017; see **Supplement**).

#### Subcortico-cortical FC Matrix Construction

For each subject, the BOLD timeseries was extracted from each of the 10 cortical networks by averaging across the vertices in a single network. The z-transformed Pearson correlation was then calculated between the BOLD timeseries of each subcortical voxel and each cortical network, resulting in a subcortical voxel x cortical network FC matrix for each subject. We excluded vertices belonging to the 10 cortical networks that were located within 10 mm of the basal ganglia or thalamus masks from the network signal in adult Talairach space (see **Supp. Methods**).

### Winner-Take-All Approach

We applied a winner-take-all approach to parcellate the basal ganglia and thalamus according to their functional relationships with cortical networks (Choi et al., 2012; Greene et al., 2014; Badke D’Andrea et al., 2023). For these analyses, FC values and matrices were first averaged across subjects. Each subcortical voxel was assigned to a single “winner” cortical network with which it correlated most. For subsequent analyses, we defined the set of voxels within either the basal ganglia or thalamus that had the same winner network as that network’s “network-selective subcortical ROI.” We assessed the reliability of the winner-take-all approach in our group-averaged data using a bootstrapping analysis (see **Supp. Methods**, **Supp. Figs. 1&2**). Group-level functional organization of the thalamus and basal ganglia was highly stable in all three ages groups, demonstrating that we had sufficient data for each age group.

**Figure 1.**
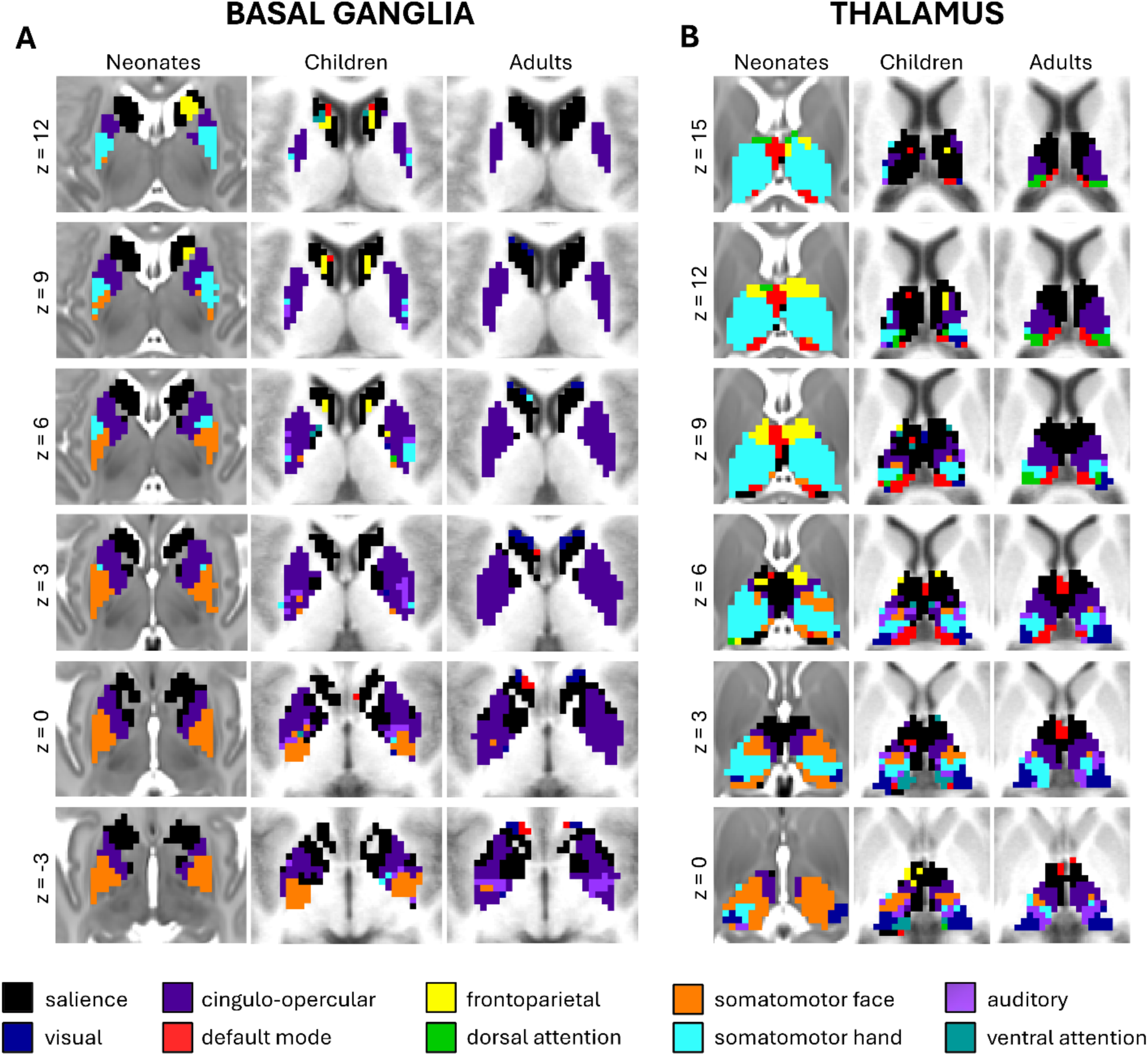
Functional network organization in A) the basal ganglia and B) the thalamus in neonates, children, and adults. Each subcortical voxel is colored according to the adult cortical network with which it has the highest FC (color key at the bottom). Slices are shown in the transverse plane (image left is anatomical left), z = -3 to z = 12 for the basal ganglia and z = 0 to z = 15 for the thalamus.

### Age Group Comparisons

To characterize how subcortico-cortical FC differed between age groups, we examined age-group differences in (1) the proportion of voxels assigned to each network, i.e., sizes of network-selective subcortical ROIs, (2) the FC strength of each network-selective ROI, and (3) the selectivity of network representations for each network-selective ROI.

#### Winner network size

Bootstrapping was used to compare the relative sizes of each network-selective subcortical ROI across age groups and determine whether these differences significantly differed from zero. For each bootstrap sample (n=1000), we generated an average winner-take-all map of the subcortex for each age group, calculated the proportion of voxels that were assigned to each of the 10 networks separately for the basal ganglia and thalamus, and then calculated the difference in these proportions between each pair of age groups. Significant differences in winner network size between age groups were determined by calculating how frequently the distribution of bootstrapped differences crossed zero.

#### Winner network strength

The strength of the FC between each network-selective ROI defined for each age group and the associated winner network was calculated for each participant in the neonatal, child, and adult groups. A two-way ANOVA with age group (neonate, child, adult), cortical network (10 networks), and group x network interaction terms evaluated how FC strength between network-selective ROIs and the cortex varied based on group and network. Post-hoc analyses consisted of network-wise, two-sample t-tests. False discovery rate was used to correct for the multiple comparisons (10 networks).

#### Winner network selectivity

The strength of the FC between each network-selective ROI defined for each age group and all other networks was also calculated for each participant in the neonatal, child, and adult groups. For each network-selective subcortical ROI, selectivity was defined as the difference between the strength of FC to the winner network and the average strength of FC to all other networks. Network selectivity was compared using a two-way ANOVA with age group (neonate, child, adult), cortical network (10 networks), and age x networks interaction as factors. To determine which networks were significantly more selective than others within each age group, we conducted one-tailed t-tests comparing the selectivity of each network-selective subcortical ROI to the average selectivity of all other network-selective subcortical ROIs separately for each age group. False discovery rate was used to correct for the multiple comparisons (10 networks).

### K-Means Clustering

In contrast with the winner-take-all approach that grouped subcortical voxels according to the FC with cortex that was strongest, k-means clustering was used to group subcortical voxels with similar patterns of FC across all cortical functional networks. Briefly, k-means clustering groups observations (here, voxels) by minimizing the Euclidean distances across multiple variables (here, FC to each of 10 networks). Clustering (k=2 through k=9) was applied to the FC between each subcortical voxel and all 10 functional networks averaged across participants, separately for the basal ganglia and thalamus and separately for each age group. For the neonatal dataset, we assessed the stability of identified clusters for k=2 through k=9 using permutation testing to determine a reasonable k value for the basal ganglia and thalamus (Lange et al., 2004; Sylvester et al., 2020; **Supp. Methods**). We concluded that k=4 was optimal for both the basal ganglia and thalamus to minimize the number of clusters while maximizing the change in stability from increasing to a greater cluster number.

## RESULTS

### The spatial layout of the functional organization of the neonatal subcortex displayed both similarities and differences to that of children and adults

Group-averaged winner-take-all maps of the basal ganglia and thalamus for neonates, children, and adults are shown in **Figure 1**. These maps illustrate, for each subcortical voxel, the cortical functional network with which it has the strongest functional connectivity (FC; Greene et al., 2014; Badke D’Andrea et al., 2023).

Qualitatively, there was a basic anterior-posterior functional organization of the subcortex across ages, such that anterior portions of the subcortex had greater representation of association networks (e.g., salience, frontoparietal), while posterior portions had greater representation of somatomotor networks. FC patterns were more similar in the basal ganglia across ages than in the thalamus. Specifically, the neonatal thalamus appeared to be almost entirely dominated by somatomotor FC (71%; **Fig. 1**), whereas this representation was smaller in children (14%) and adults (10%). These trends were also evident when treating each cortical network as a seed and examining the pattern of FC within the subcortex (**Supp. Fig. 3**). Similar functional organization was observed in neonates when dividing the subcortex using cortical functional networks defined in neonates (Myers et al., 2024) or a network solution that differentiates the somatomotor hand and leg networks (Gordon et al., 2017).

### Network-selective subcortical regions differ in their size, functional connectivity strength to cortex, and network selectivity across neonates, children, and adults

We next characterized how each identified region of interest (ROI) with preferential FC to a specific cortical network (“network-selective subcortical ROIs”) differed across neonates, children, and adults.

#### Age-related differences in the relative sizes of network-selective subcortical ROIs

The relative sizes of each network-selective subcortical ROI within each age group are shown in **Figure 2A & C** and **Tables 1 & 2** and the differences between age groups are displayed in **Figure 2B & D** (neonates vs. children), **Supplementary Figure 6** (neonates vs. adults, children vs. adults) and **Tables 1 & 2**. All somatomotor network-selective subcortical ROIs were larger (+7.8%-+40.4%) in neonates than children for both the basal ganglia (**Table 1**, **Fig. 2B**) and thalamus (**Table 2**, **Fig. 2D**). Consistent with previous reports (Greene et al., 2014; Badke D’Andrea et al., 2023), somatomotor face network-selective subcortical ROIs were larger in children than in adults (**Tables 1 & 2**, **Supp. Fig. 6B & D**), demonstrating a consistent decrease in the size of somatomotor representation in the subcortex over the course of development. Conversely, many association network-selective subcortical ROIs (salience, cingulo-opercular, default mode) were larger (+4.7%-+33%) in older cohorts than in neonates. One notable exception was the frontoparietal network, which was larger in neonates than in older cohorts (**Tables 1 & 2**, **Fig. 2B & D**, **Supp. Fig. 6**). Further, some functional networks had no neonatal representation in the subcortex and thus were by definition larger in older cohorts than in the neonates (e.g., visual and auditory in the basal ganglia, ventral attention in the thalamus; **Tables 1 & 2**, **Fig. 2**, **Supp. Fig. 6**).

**Figure 2.**
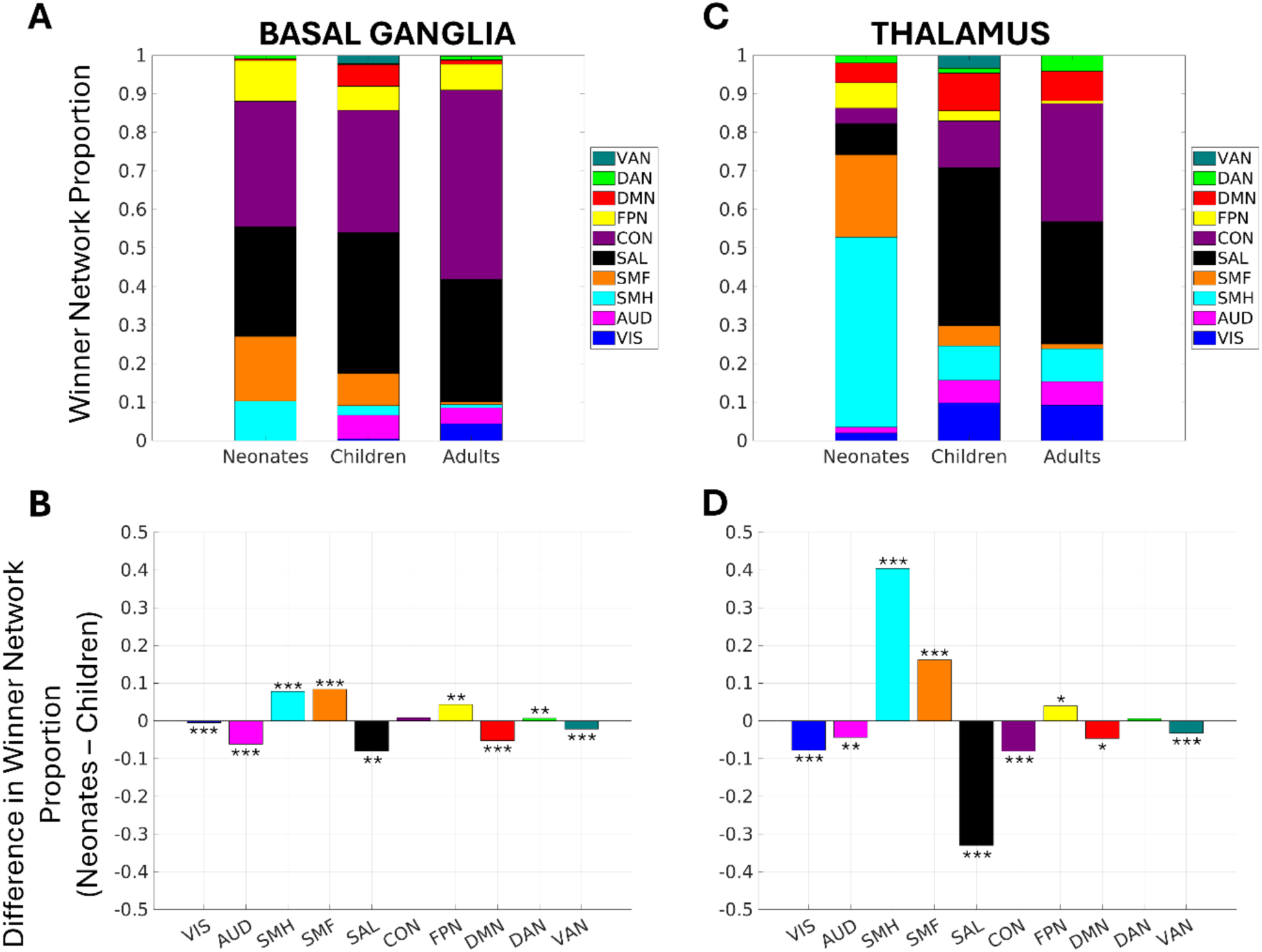
Age-related differences in the relative sizes of network-selective subcortical ROIs. **A, C)** Stacked bar graphs illustrating the relative proportion of **A)** the basal ganglia and **C)** the thalamus that each network represents according to the winner-take-all results. **B, D)** Bar graphs showing the difference in the relative size of network-selective ROIs in **B)** the basal ganglia and **D)** the thalamus between neonates and children. A bar above the x-axis indicates that the network-selective ROI is larger in neonates than in children. A bar below the x-axis indicates that the network-selective ROI is smaller in neonates than in children. Difference bar graphs comparing neonates vs. adults and children vs. adults are shown in **Supplementary** Figure 6. Significant differences in relative size between ages were determined via bootstrapping. *p<0.05, **p<0.01, ***p<0.001.

**Table 1.** Relative sizes of network-selective ROIs in the basal ganglia and statistical significance of age-related differences. Plus (+) signs refer to larger proportions in the older cohorts compared to younger cohorts. Minus (-) signs refer to smaller proportions in the older cohorts compared to younger cohorts.

|  | Proportion of each network-selective basal ganglia ROI |  |  | Age comparison statistics |  |  |
| --- | --- | --- | --- | --- | --- | --- |
|  | Neonates | Children | Adults | Neonates v. Children | Neonates v. Adults | Children v. Adults |
| **VIS** | 0.0% | 0.5% | 4.4% | +0.5%, p<0.001 | +4.4%, p<0.001 | +3.9%, p<0.001 |
| **AUD** | 0.0% | 6.2% | 4.1% | +6.2%, p<0.001 | +4.1%, p<0.001 | -2.1%, n.s. |
| **SMH** | 10.3% | 2.5% | 0.8% | -7.8%, p<0.001 | -9.5%, p<0.001 | -1.7%, n.s. |
| **SMF** | 16.8% | 8.4% | 0.7% | -8.4%, p<0.001 | -16.1%, p<0.001 | -7.7%, p<0.001 |
| **SAL** | 28.5% | 36.5% | 31.8% | +8.0%, p<0.01 | +3.3%, n.s. | -4.7%, n.s. |
| **CON** | 32.6% | 31.8% | 49.1% | -0.8%, n.s. | +16.5%, p<0.001 | +17.3%, p<0.001 |
| **FPN** | 10.6% | 6.2% | 6.7% | -4.4%, p<0.01 | -3.9%, p<0.001 | +0.5%, n.s. |
| **DMN** | 0.4% | 5.7% | 1.1% | +5.3%, p<0.001 | +0.7%, n.s. | -4.6%, p<0.001 |
| **DAN** | 0.9% | 0.2% | 1.0% | -0.7%, p<0.001 | +0.1%, n.s. | +0.8%, n.s. |
| **VAN** | 0.0% | 2.2% | 0.2% | +2.2%, p<0.001 | +0.2%, p<0.001 | -2.0%, p<0.05 |

**Table 2.** Relative sizes of network-selective ROIs in the thalamus and statistical significance of age-related differences. Plus (+) signs refer to larger proportions in the older cohorts compared to younger cohorts. Minus (-) signs refer to smaller proportions in the older cohorts compared to younger cohorts.

|  | Proportion of each network-selective thalamic ROI |  |  | Age comparison statistics |  |  |
| --- | --- | --- | --- | --- | --- | --- |
|  | Neonates | Children | Adults | Neonates v. Children | Neonates v. Adults | Children v. Adults |
| **VIS** | 2.0% | 9.8% | 9.3% | +7.8%, p<0.001 | +7.3%, p<0.001 | -0.5%, n.s. |
| **AUD** | 1.5% | 5.9% | 6.1% | +4.4%, p<0.01 | +4.6%, n.s. | +0.2%, n.s. |
| **SMH** | 49.3% | 8.9% | 8.5% | -40.4%, p<0.001 | -40.8%, p<0.001 | -0.4%, n.s. |
| **SMF** | 21.4% | 5.3% | 1.2% | -16.1%, p<0.001 | -20.2%, p<0.001 | -4.1%, p<0.001 |
| **SAL** | 8.0% | 41.0% | 31.8% | +33.0%, p<0.001 | +23.8%, p<0.001 | -9.2%, p<0.05 |
| **CON** | 4.1% | 12.1% | 30.7% | +8.0%, p<0.001 | +26.6%, p<0.001 | +18.6%, p<0.001 |
| **FPN** | 6.6% | 2.6% | 0.7% | -4.0%, p<0.05 | -5.9%, p<0.001 | -1.9%, n.s. |
| **DMN** | 5.1% | 9.8% | 7.7% | +4.7%, p<0.05 | +2.6%, n.s. | -2.1%, n.s. |
| **DAN** | 1.9% | 1.3% | 4.1% | -0.6%, n.s. | +2.2%, n.s. | +2.8%, p<0.05 |
| **VAN** | 0.1% | 3.3% | 0.0% | +3.2%, p<0.001 | -0.1%, n.s. | -3.3%, p<0.001 |

#### Age-related differences in the functional connectivity strength of network-selective subcortical ROIs

Winner network strength for each network-selective subcortical ROI in neonates, children, and adults is shown in **Figure 3**. Applying a two-way ANOVA to the winner network strength for each network-selective subcortical ROI in the neonates, children, and adults revealed a significant main effect of age group (BG: f=8.56, p<0.001; THAL: f=133.66, p<0.001) and of network (BG: f=4.27, p<0.001; THAL: f=27.36, p<0.001) for both the basal ganglia and thalamus. There was also a significant age group x network interaction for both the basal ganglia (f=27.53, p<0.001) and the thalamus (f=5.60, p<0.001), indicating different developmental trajectories for different networks. Post-hoc analyses compared the winner network strength for each network-selective subcortical ROI between age groups separately for the basal ganglia (**Fig. 3A**) and thalamus (**Fig. 3B**) to further interrogate these effects.

**Figure 3.**
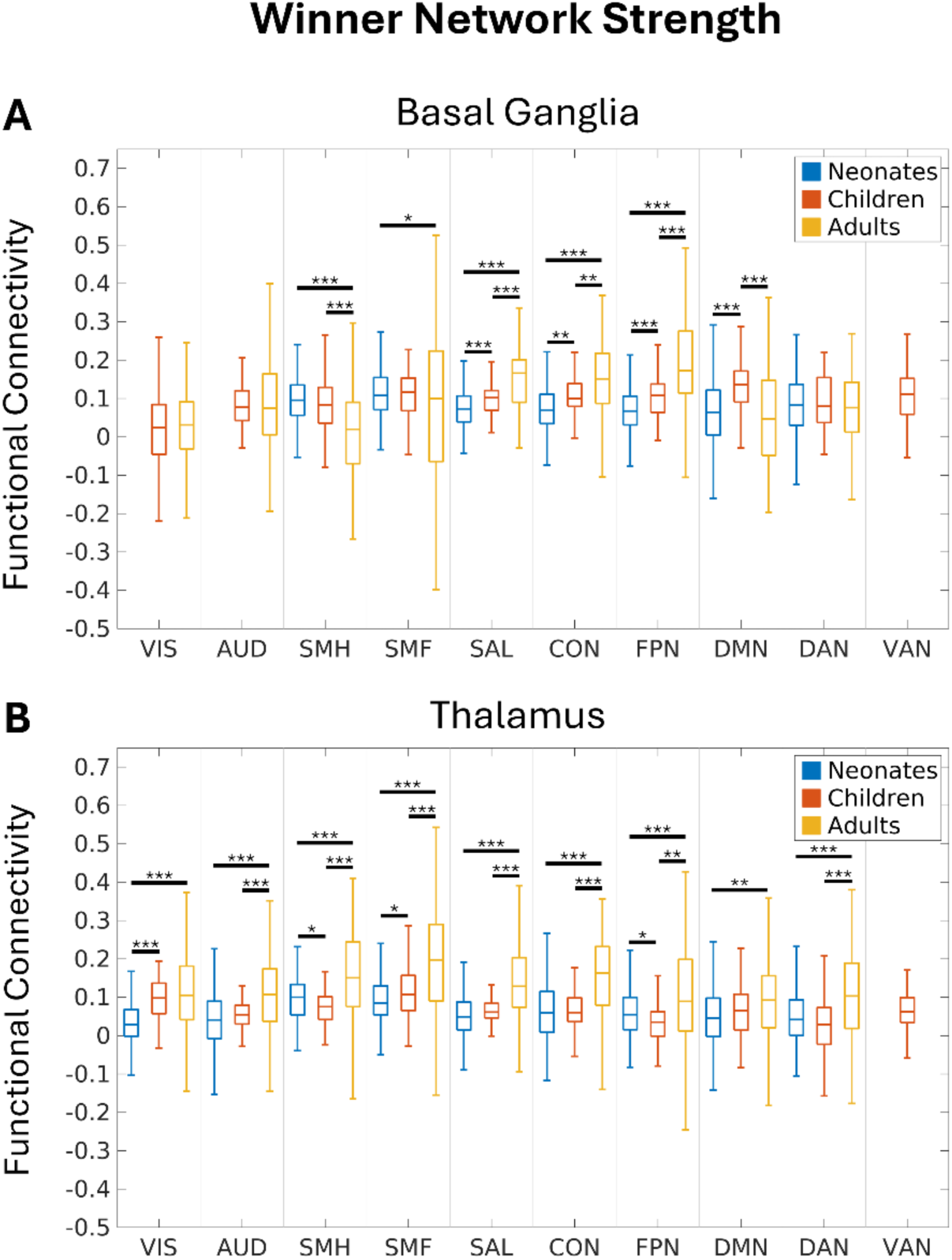
Age-related differences in FC strength of network-selective subcortical ROIs in A) the basal ganglia and B) the thalamus. Box-and-whisker plots of network-wise FC strength for neonates (blue), children (red), and adults (yellow), organized by network. Statistical significance of FC strength between ages for each network was determined by two-tailed t-tests. *p<0.05, **p<0.01, ***p<0.001.

In the basal ganglia, developmental patterns of winner network strength differed for somatomotor and association networks. Winner network strength for the somatomotor hand network-selective ROI was significantly higher in neonates (t=9.05, p<0.001) and children (t=4.07, p<0.001) than in adults. The somatomotor face network-selective ROI also exhibited higher winner network strength in neonates compared to adults (t=2.58, p=0.014). In contrast, several association network-selective ROIs exhibited higher FC strength in older cohorts than in younger cohorts, including the frontoparietal (neonates vs. adults: t=-11.92, p<0.001; children vs. adults: t=-4.94, p<0.001), salience (neonates vs. children: t=-4.05, p<0.001, neonates vs. adults: t=-11.28, p<0.001; children vs. adults: t=-4.76, p<0.001), and cingulo-opercular (neonates vs. children: t=-3.43, p<0.01; neonates vs. adults: t=-8.44, p<0.001; children vs. adults: t=-3.26 p<0.01) network-selective ROIs.

In the thalamus, there was a shared developmental pattern such that winner network strength generally increased with age for each network-selective subcortical ROI. Winner network strength was significantly greater (p<0.01) in adults than in neonates for all network-selective subcortical ROIs and significantly greater in adults than in children for 8 of the 10 (except the default mode and visual networks). While winner network strength tended to be greater in older cohorts, winner network strength did not significantly differ between neonates and children for most network-selective subcortical ROIs, and for the somatomotor hand network-selective ROI, winner network strength was significantly weaker in children compared to neonates (t=2.59, p=0.0225).

#### Age-related differences in winner network selectivity of network-selective subcortical regions

Age-related differences in winner network selectivity are depicted in **Figure 4**. Applying a two-factor ANOVA to the winner network selectivity (i.e., FC to winner network – average FC to other networks) for each network-selective subcortical ROI in the neonates, children, and adults revealed a significant main effect of age group (BG: f=59.6, p<0.001; THAL: f=343.5, p<0.001) and of network (BG: f=3.75, p=0.001; THAL: f=15.07, p<0.001). There was also a significant age group x network interaction for both the basal ganglia (f=30.91, p<0.001) and the thalamus (f=6.10, p<0.001), indicating different developmental trajectories for different networks. Post-hoc one-tailed t-tests identified which network-selective subcortical ROIs exhibited significantly greater winner network selectivity than other networks separately for each age group to further interrogate this interaction. Post-hoc comparisons of winner network selectivity between neonates, children, and adults for each network-selective subcortical ROI are listed in **Supplementary Table 1** and indicated a general trend of increased selectivity of network-selective subcortical ROIs with age.

**Figure 4.**
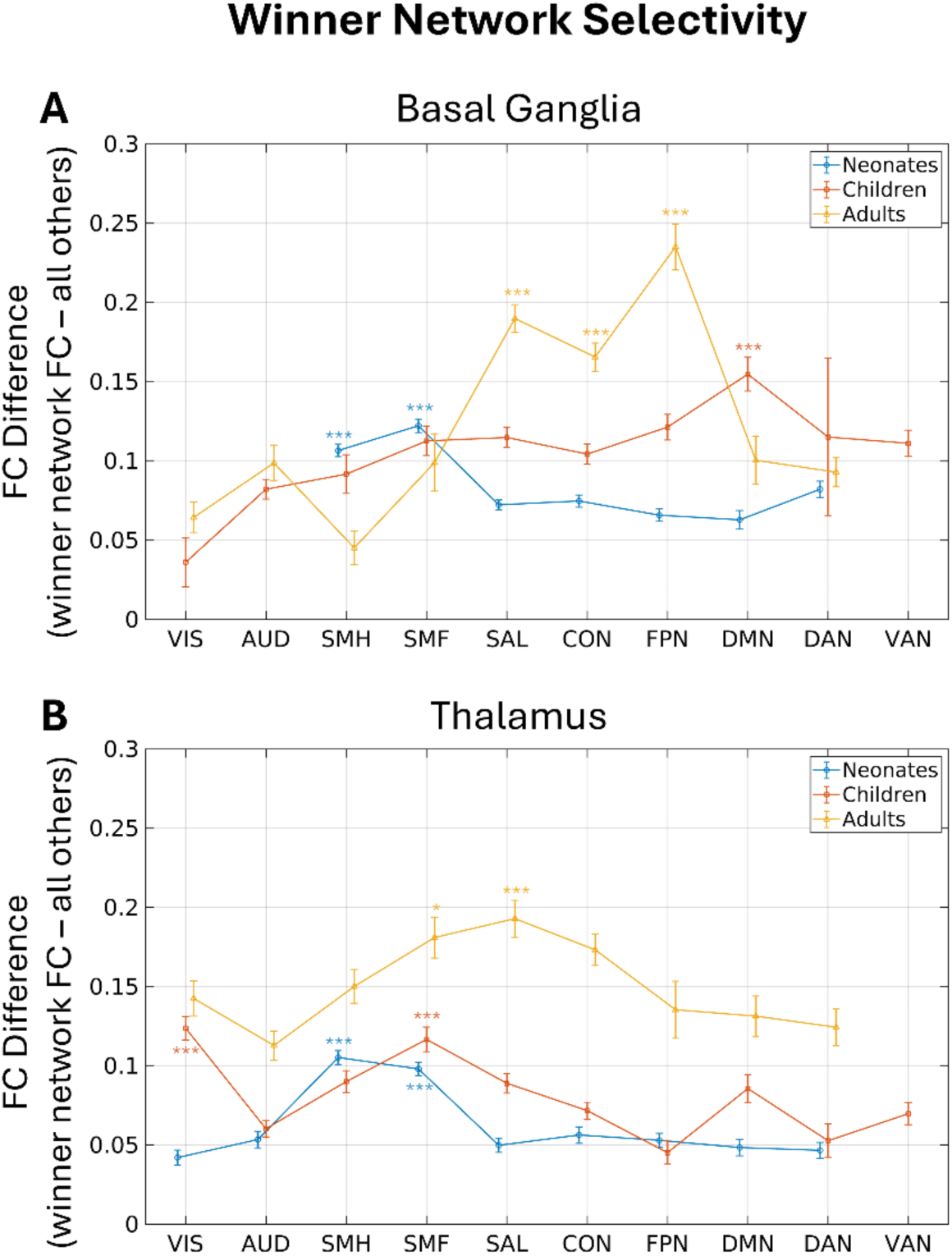
Age-related differences in winner network selectivity of network-selective subcortical ROIs in A) the basal ganglia and B) the thalamus. Line plots of winner network selectivity for neonates (blue), children (red), and adults (yellow). Selectivity was defined as the difference between the FC of the winner network and the average FC of all other networks. Error bars display standard error of the mean. Statistical significance of winner network selectivity within each age group was determined by one-tailed t-tests. *p<0.05, **p<0.01, ***p<0.001.

For both the basal ganglia and thalamus, which network-selective subcortical ROIs that exhibited the greatest winner network selectivity differed by age group. In neonates (**Fig. 4**, blue), winner network selectivity was greatest for network-selective subcortical ROIs representing somatomotor networks. Subcortical representations of the somatomotor hand (BG: t=5.46, p<0.001; THAL: t=9.58, p<0.001) and the somatomotor face (BG: t=9.34, p<0.001, THAL: t=7.95, p<0.001) networks in both the basal ganglia and thalamus exhibited significantly more selective FC to the cortex than other network-selective subcortical ROIs. In children (**Fig. 4**, red), winner network selectivity was greatest for representations of the default mode network (t=5.34, p<0.001) in the basal ganglia and the visual (t=5.83, p<0.001) and somatomotor face (t=4.83, p<0.001) networks in the thalamus. In adults (**Fig. 4**, yellow), higher winner network selectivity was observed in the network-selective subcortical ROIs representing the frontoparietal (t=9.52, p<0.001), salience (t=5.59, p<0.001) and cingulo-opercular (t=3.58, p<0.001) networks in the basal ganglia and the salience (t=3.75, p<0.001) and somatomotor face (t=2.71, p=0.0153) networks in the thalamus. These results suggest that subcortical representations of some sensorimotor systems tend to be more selective in younger ages, while subcortical representations of some association networks are more selective in adults.

### Clusters within the neonatal basal ganglia and thalamus display patterned integration of networks that ranged from association- to sensorimotor-dominant

While straightforward, the winner-take-all approach does not incorporate any secondary but biologically significant functional connections to other cortical networks that may exist in the subcortex. For all age groups, many of the network-selective subcortical ROIs described above exhibited strong FC with multiple other cortical functional networks (**Supp. Figs. 7-9**). Thus, we employed a data-driven clustering approach to group voxels with similar patterns of subcortical-cortical FC to more directly interrogate functional network integration in the neonatal basal ganglia and thalamus. **Figure 5** displays the clusters resulting from k-means clustering (k=4) and the average FC between each cluster and each functional network for both the neonatal basal ganglia and the thalamus. K-means clustering was performed for k=3 through k=7 (**Supp. Figs. 10-18)** and stability analyses indicated that k=4 was a suitable choice for both the basal ganglia and the thalamus (**Supp. Fig. 19**).

**Figure 5.**
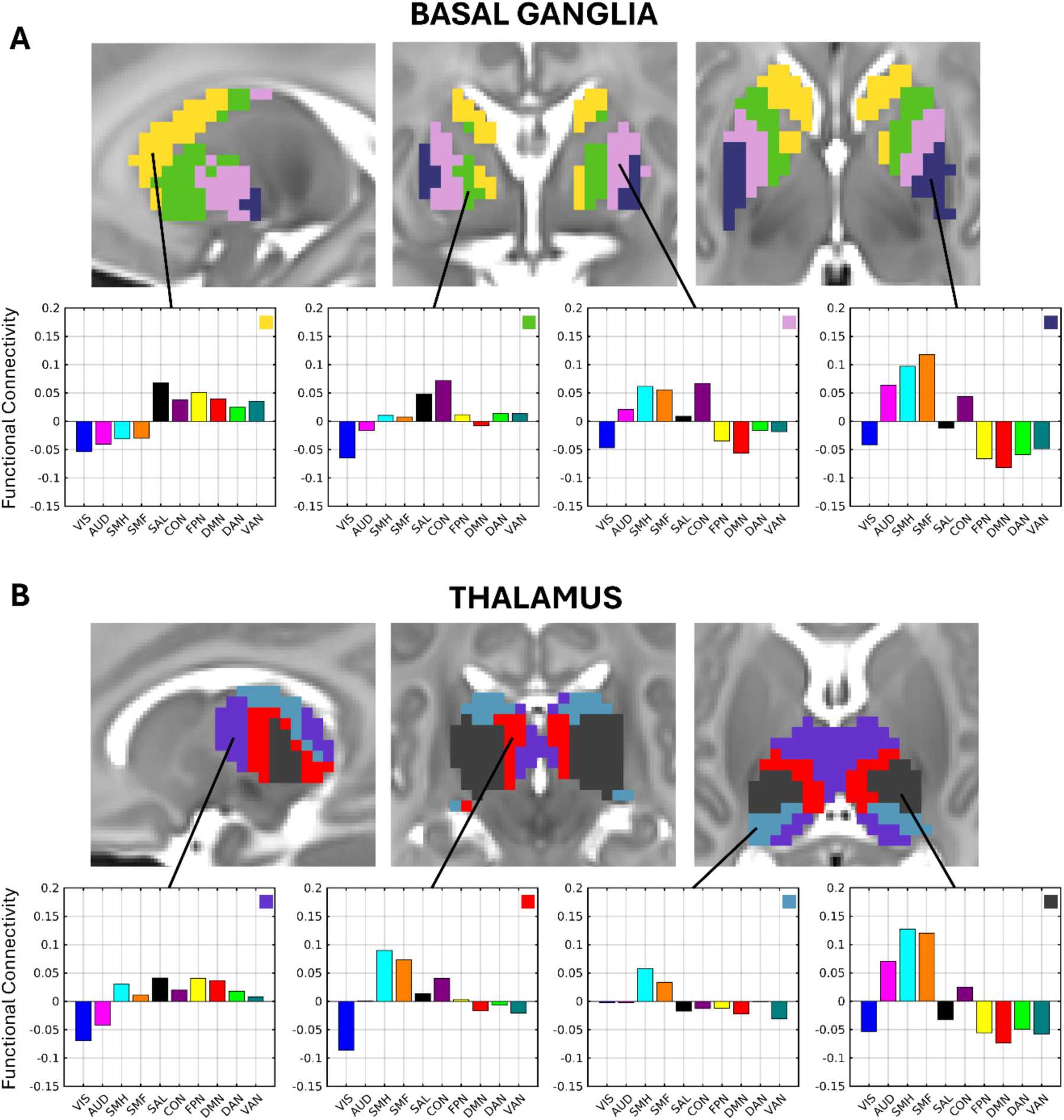
K-means clusters within the neonatal. **A**) basal ganglia (gold, green, pink, indigo) and **B)** thalamus (purple, red, blue, dark gray) according to overall network profiles. Sagittal (left), coronal (middle), and transverse (right) slices are displayed. Basal ganglia coordinates: x = -20, y = -3, z = 3. Thalamus coordinates: x = -12, y = -24, z = 9. Average network profiles showing average FC to each network for the voxels within each cluster are depicted below. Each network profile graph contains a box with its corresponding cluster color in the top right.

In both the neonatal basal ganglia and thalamus, there was a clear transition from more association-network-dominant (e.g., **Fig. 5A**, gold cluster; **Fig. 5B**, purple cluster) to sensorimotor-network-dominant clusters (e.g., **Fig. 5A**, indigo cluster; **Fig. 5B**, dark gray) along the anterior-posterior axis. In the basal ganglia, the anterior gold and green clusters exhibited strong FC with only association networks (gold: salience, cingulo-opercular, frontoparietal, default mode, dorsal attention, ventral attention; green: salience, cingulo-opercular) while the posterior indigo cluster exhibited strong FC with largely sensorimotor networks (indigo: somatomotor hand, somatomotor face, auditory). In the thalamus, each cluster exhibited positive FC with the somatomotor hand network, but the FC to other networks differed such that the anterior purple cluster also exhibited strong FC to association networks (purple: salience, frontoparietal, default mode) and the posterior dark gray and teal clusters exhibited strong FC to other sensorimotor networks (dark gray: somatomotor face, auditory; teal: somatomotor face). Interleaved between these sensorimotor- or association-dominant clusters in both the basal ganglia and thalamus were clusters that exhibited strong FC to the cingulo-opercular network and somatomotor face and hand networks (e.g., **Fig. 5A**, pink cluster; **Fig. 5B**, red cluster), demonstrating subcortical integration across sensorimotor and association networks.

K-means clustering was also applied to the subcortical-cortical FC from the children and adult datasets in **Supplementary Figures 20-29** for qualitative comparison. Consistent with the observed age-related differences in winner network selectivity described above, the FC of the clusters identified in children and adults appeared to exhibit greater network selectivity (i.e., less integration) than those identified in the neonates (**Supp. Figs. 20-29**).

## DISCUSSION

The goals of this study were to map the functional organization of the neonatal basal ganglia and thalamus and to characterize key features of subcortico-cortical FC (size, strength, and selectivity of network representations) in neonates and how they differ in children and adults. The overall functional organization of the neonatal basal ganglia largely aligns with that of children and adults, with anterior portions exhibiting greater association network representation and posterior portions exhibiting greater sensorimotor network representation. The neonatal thalamus contains some basic anterior-posterior organization as well but diverges from children and adults in its overwhelming somatomotor representation. In both the basal ganglia and thalamus, there were significant differences in the size, strength, and selectivity of network representations between neonates, children, and adults. The relative size of the somatomotor representation in the subcortex was greater in neonates compared to children and adults. Further, the strength and selectivity of FC between the subcortex and somatomotor networks tended to be higher in neonates than in older cohorts, whereas the strength and selectivity of FC between the subcortex and association networks were generally higher in children and adults. Finally, the clusters identified in the neonatal subcortex with similar patterns of FC to all functional networks corroborated the anterior-posterior organization of association-sensorimotor network representations but also provided evidence of integration as multiple functional networks were represented in each cluster.

### Functional organization can be observed in the subcortex within the first month of life

Our findings indicate that the representation of multiple functional networks in the subcortex and the overarching anterior-posterior spatial layout of association versus sensorimotor functional representations present in children and adults (Greene et al., 2014; Badke D’Andrea et al., 2023) is already present shortly after birth in the basal ganglia. The somatomotor overrepresentation that we found in the thalamus aligns with prior work (Alcauter et al., 2014; Toulmin et al., 2015), despite differences in our methodology (e.g., higher amounts of low-motion data, pre-processing that incorporates the individual neonatal cortical surface, dividing the cortex using functional network templates). Prenatal developmental processes (e.g., subcortical-to-cortical axonal outgrowth and spontaneous wave-like propagation) likely establish these distinctions within the subcortex that support the foundations of emerging cortico-striato-thalamo-cortical loops (Kerschensteiner, 2014; Martini et al., 2021; Klavinskis-Whiting et al., 2023; Oldham et al., 2025).

### A relative shift in sensorimotor versus association functional network representation in the subcortex over development

Somatomotor network-selective subcortical ROIs comprised a significantly larger proportion of subcortical volumes in neonates than in children or adults. Non-linear expansion of the basal ganglia and thalamus from birth to childhood is one possible explanation, such that the FC of each individual subcortical nucleus is stable over development, but the volume of nuclei representing association networks expand more than somatomotor nuclei. Consistent with this hypothesis, prior work using structural and diffusion MRI has demonstrated that sensorimotor thalamic nuclei expand and form earlier than association nuclei in the third trimester and early postnatal period (Zheng et al., 2023) and that thalamic volume increases rapidly from the anterior portion rather than homogeneously during the first few postnatal years (Tutunji et al., 2018). Alternatively, the overrepresentation of somatomotor functional networks in the neonatal subcortex may reflect a reorganization of subcortico-cortical functional connectivity such that nuclei are associated with different functional networks over different stages of postnatal development. Synaptogenesis and pruning may alter the relative weights of regional subcortical connections to different portions of cortex, such that some areas of the subcortex may shift from being more closely linked to sensorimotor networks to being more closely linked association networks. Consistent with reorganization, there is substantial evidence that subcortico-cortical connections mature earlier between motor cortices than others via increased synaptogenesis, myelination, and maturation of inhibition (Grant et al., 2012; Knowles et al., 2021; Sydnor et al., 2021, 2025). It is possible that both non-linear expansion and functional reorganization contribute to the observed overrepresentation of somatomotor networks in the neonatal subcortex when compared to older children and adults.

The strength of FC between somatomotor networks and the subcortex was also generally greater in neonates than in older cohorts (except the child thalamus) and the strength of FC between association networks and the subcortex was weaker in neonates than in older cohorts. FC strength is thought to reflect the statistical history of co-activation achieved through Hebbian learning (Fox et al., 2005), and as such, the observed variation in FC strength between the subcortex and different functional networks across age groups may reflect developmental differences in the co-recruitment of the cortex and subcortex during daily life. Emerging neurotransmitter function in the globus pallidus (Valenti et al., 2003), the location of somatomotor network-selective ROIs in the basal ganglia, may also explain the observed differences in FC strength with age. During early postnatal development, GABAergic synaptic activity transitions from being excitatory to inhibitory (Ganguly et al., 2001; Lozovaya et al., 2023), potentially contributing to the isolated decreases in FC between the basal ganglia and somatomotor systems after the neonatal timepoint. Axonal pruning and learning (Cowan et al., 1984) that occur later in development may also explain the observed decreased FC strength between the subcortex and somatomotor cortex. Prior studies are consistent with these developmental trends beginning at birth, demonstrating a continuous decrease in FC strength of the somatomotor face network representation in both the basal ganglia and the thalamus through childhood and adolescence (Greene et al., 2014; Badke D’Andrea et al., 2023).

Consistent with studies of cortical functional networks in neonates, children, and adults (Badke D’Andrea et al., 2023; Sylvester et al., 2023; Tooley et al., 2024), the selectivity of the FC between network-selective subcortical ROIs and the cortex increased with age. Prenatal synaptic pruning and refinement and postnatal experience-dependent mechanisms likely sculpt and tune subcortical-cortical FC to increase network selectivity (Gilmore et al., 2018; Nielsen et al., 2023; Sydnor et al., 2023, 2025; Zheng et al., 2023). The relative network selectivity of different subcortical representations differed per age group such that somatomotor representations in the neonatal subcortex exhibited greater selectivity than association representations, but the opposite was true for children and adults. Thus, the mechanisms associated with the refinement of subcortical-cortical FC likely do not occur at the same rate, with subcortical representations of sensorimotor networks developing earlier than representations of association networks.

Overall, our results support our original hypothesis that the development of the functional organization of the subcortex reflects the pattern of neurodevelopment that occurs across the cortex, with sensorimotor systems generally developing earlier than association systems. In the cortex, the FC of sensorimotor networks exhibit greater maturity (i.e., stronger within-network FC, greater network selectivity) relative to other networks at birth, and we find this same developmental trend in the neonatal subcortex. Early maturation of the representations of sensorimotor networks in the subcortex may have behavioral consequences for the neonate. It is possible that the neonatal subcortex is primed to process somatomotor information through early experiences in utero. Fetal movements and reflexes are associated with the propagation of feedforward and feedback activity that travels between the spinal cord, subcortex, and cortex which tune developing somatosensory and motor circuits and shape emerging behavior (Stockx et al., 2007). Typical development of motor skills early in life is important for subsequent cognitive development because infants need to physically explore their environment to learn and develop cognitive concepts (Piaget, 1952; Murray et al., 2007; Shi and Feng, 2022). The observed differences in the developmental trajectories of sensorimotor and association network representations in the subcortex may reflect this developmental cascade of emerging brain function. As an individual grows and develops motor capabilities, subcortico-cortical loops can begin to prioritize higher order information processing; thus, association representations in the subcortex become larger, stronger, and more selective with age.

### The clustering of multiple functional networks within the neonatal subcortex may provide early evidence of motor integration zones

Using a data-driven approach that allows the representation of more than one functional network, we additionally identified clusters in the neonatal basal ganglia and thalamus composed of either multiple sensorimotor networks, multiple association networks, or a mix of sensorimotor and association networks. These clusters did not exhibit strong selectivity for a single functional network. There are prominent secondary connections throughout the neonatal subcortex that are not revealed through a winner-take-all approach. One explanation for this lack of functional specialization in the neonatal subcortex may be the immaturity and weak differentiation of neonatal cortical functional networks (Gao et al., 2017; Grayson and Fair, 2017; Sylvester et al., 2023). As cortical network segregation increases over the course of development (Tooley et al., 2024), representations of a single functional network within the subcortex may become more well-defined. Consistent with this explanation, the same data-driven clustering of the subcortex revealed less network integration in the children and adults (**Supp. Figs. 20-29**).

Simultaneous representation of multiple functional networks in the neonates might reflect integration across cortico-striato-thalamo-cortical loops. For example, we identified clusters in the neonatal basal ganglia and thalamus that had relatively strong FC to both somatomotor networks and the cingulo-opercular network. The cingulo-opercular network, also called the “action-mode network” because of its frequent coordination during motor behavior (Dosenbach et al., 2025), is involved in executive control of task performance (Dosenbach et al., 2007; Sadaghiani and D’Esposito, 2015; Crittenden et al., 2016). In children and adults, integrated representation of both somatomotor and cingulo-opercular networks has been observed in select portions of the subcortex, termed the “motor integration zone” (Greene et al., 2020; Badke D’Andrea et al., 2023) and is thought to be involved in top-down control of motor systems. The presence of early motor integration zones in the neonatal subcortex may indicate that the development of executive control over movement is an early priority for human brain development, as infants gradually learn to control basic goal-directed movements. More broadly, the profile of subcortical-cortical FC may indicate which functional networks are already starting to work together through integration in the cortico-striato-thalamo-cortical loops beginning at birth.

### Clinical significance of altered subcortical functional organization in neonates

Importantly, the development of the key features of subcortico-cortical FC described in this work (size, strength, and selectivity of network representations) have been found to be altered in several developmental disorders (Nair et al., 2013; Cerliani et al., 2015; Brown et al., 2017; Abbott et al., 2018; Ramkiran et al., 2019). For example, in children with Tourette syndrome, the size and strength of the representations of sensorimotor networks in the subcortex are increased and the selectivity of the FC between these sensorimotor representations and other sensorimotor and association networks is decreased when compared to healthy controls (Ramkiran et al., 2019). Because functional connectivity tracks the statistical history of co-activation across the lifespan, these alterations may contribute to the frequency/severity of tics and other symptoms and/or may arise as a consequence of experiencing and managing tics (Greene et al., 2025). Understanding whether these features are altered during infancy before tics or other symptoms emerge would disambiguate these explanations and potentially provide useful targets for treatment and intervention. Thus, our work characterizing how the features of subcortical functional organization emerge during normative development beginning at birth is critical to future investigations of how differences in subcortical functional organization may give rise to disordered development.

### Limitations

Comparisons between neonates, children, and adults are complicated by multiple unavoidable factors. Differences in the imaging parameters and data processing of each dataset are necessary to handle the differences in head size, magnetic properties of tissues, and sources of noise (Thornton et al., 1999; Smyser and Neil, 2015; Liu et al., 2016; Goksan et al., 2017; Neil and Smyser, 2018; Kaplan et al., 2022). While different scanner sequences should have little impact on the relative size and relative strength of connections, direct comparison of the absolute strength of FC between neonates and older cohorts likely reflects a combination of methodological and biological differences. Second, all neonatal data were collected during natural sleep, while children and adult data were collected during resting wake. Sleep state impacts some features of FC in adults (Tagliazucchi and Laufs, 2014), but its impact on features of neonatal FC is less clear (Korom et al., 2022). Furthermore, we used the same adult network template (Gordon et al., 2016) in winner-take-all analyses for all ages to facilitate age comparisons even though these functional network templates may not best capture neonatal FC (Tu et al., 2025). However, the observed functional organization of neonatal subcortex was largely replicated when using a neonatal-specific functional network parcellation (Myers et al., 2024). Finally, group-averaging and cross-sectional analyses obscure individual variation and within-subject longitudinal changes in subcortical functional organization. Since individual differences in this organization may be clinically relevant in adults (Greene et al., 2020), future work should investigate the extent to which individual variation in subcortical functional organization is present at birth and how these features change with age within individuals in order to better understand disordered development (Labonte et al., 2024, 2025).

### Conclusions

The current study demonstrates that the basic anterior-posterior organization of association to sensorimotor representation in the basal ganglia and thalamus is established at birth, suggesting that this organization emerges prenatally. Features of subcortico-cortical FC largely evolve along the sensorimotor-association axis of development, with sensorimotor areas having relatively larger representations and stronger and more selective FC to cortical networks. This somatomotor dominance at birth may reflect current experiences of the neonate (e.g. sensory/motor experiences) and/or the need for continued development to refine the representation and integration of information from multiple networks. This characterization of the state of subcortico-cortical functional organization in healthy, full-terms at birth serves as a starting point for tracking normative development versus development linked to adverse prenatal exposures, premature delivery, and/or psychiatric disorders.

## Acknowledgements

The authors thank the families of all our participants from these studies for their participation and dedication toward research.

## CRediT Authorship Contribution Statement

**Samantha L. Blake**: Conceptualization, Methodology, Formal analysis, Visualization, Writing – Original Draft, Writing – Review & Editing; **Jeanette K. Kenley**: Data curation, Writing – Review & Editing; **Tara A. Smyser**: Data curation, Writing – Review & Editing; **Aidan Latham**: Data curation; **Dimitrios Alexopoulos**: Data curation; **Deanna J. Greene**: Writing – Review & Editing; **Rachel E. Lean**: Writing – Review & Editing; **Deanna M. Barch**: Writing – Review & Editing, Funding acquisition, Resources; **Barbara B. Warner**: Writing – Review & Editing, Funding acquisition, Resources; **Joan L. Luby**: Writing – Review & Editing, Funding acquisition, Resources; **Cynthia E. Rogers**: Writing – Review & Editing, Funding acquisition, Resources; **Christopher D. Smyser**: Writing – Review & Editing, Funding acquisition, Resources, Supervision; **Chad M. Sylvester**: Conceptualization, Writing – Original Draft, Writing – Review & Editing, Supervision; **Ashley N. Nielsen**: Conceptualization, Methodology, Writing – Original Draft, Writing – Review & Editing, Supervision.

