## Supplemental Material for "Functional Organization of the Neonatal Basal Ganglia and Thalamus"

**SUPPLEMENTARY METHODS**

**Participants**

*Inclusion and exclusion criteria*

Inclusion and exclusion criteria were similar across the three datasets. Inclusion criteria for neonates included maternal age >18 years and full-term singleton birth (>37 weeks gestational age). Excluded were mothers with alcohol or other substance abuse, subjects with anatomical abnormalities or injuries based on structural MRI scans, women with pregnancy complications, and subjects with known fetal abnormalities.

All child and adult participants were English-speaking, were right-handed, and reported no history of neurological or psychiatric disease.

**Image Acquisition: Neonates**

Before scanning, neonates were fed, swaddled, and positioned in a head-stabilizing vacuum fix wrap (Mathur et al. 2008). A nurse qualified in neonate transport and resuscitation was present at all MRI scans. Heart rate and blood oxygenation were measured continuously throughout all scans, and infants were monitored via video. We acquired between 2 and 9 fMRI BOLD scans depending on infant tolerability (mean 3.75 runs). Runs were collected in both the anterior-to-posterior (AP) and posterior-to-anterior (PA) directions, with a typical scan session consisting of 2 AP runs and 2 PA runs. AP and PA scans were concatenated following fMRI preprocessing but before FC processing.

**Image Acquisition: Children**

Imaging for the child dataset was performed on a Siemens MAGNETOM PRISMA 3T MRI Scanner with a 32-channel head coil. During scanning, children were instructed to watch a centrally presented white crosshair on a black background, and to relax and remain as still as possible. T1-weighted sagittal MP-RAGE (magnetization-prepared rapid acquisition gradient echo) structural images (TE=2.9 ms, TR=2500 ms, inversion time (TI)=1090 ms, flip angle=8°, 176 slices, 1-mm isotropic resolution) and T2-weighted images (sagittal, 176 slices, 1-mm isotropic resolution, TE=565 ms, TR=3200 ms) were collected. Functional MRI was performed using a BOLD contrast sensitive gradient echo-planar sequence (TE=30 ms, TR=800 ms, flip angle=52°, 2.4-mm isotropic resolution, 60 axial slices, MB factor=6). The number of collected volumes ranged from 561 to 2901 (mean = 1520 frames and 20.3 min). 2 AP and 2 PA field maps were obtained during each session with the same parameters.

**Image Acquisition: Adults**

Imaging for the adult dataset was performed on a Siemens MAGNETOM Trio Tim 3T Scanner with a Siemens 12-channel Head Matrix Coil. Adults were also instructed to watch a central white crosshair and remain still. T1-weighted sagittal MP-RAGE structural images (TE=3.06 ms, TR=2400 ms, TI=1 s, flip angle=8°, 127 slices, 1-mm isotropic resolution) and T2-weighted turbo spin echo structural images (TE=84 ms, TR=6.8 s, 32 slices, 2 x 1 x 4-mm voxels) were collected. Functional MRI was performed using a BOLD contrast sensitive gradient echo-planar sequence (TE=27 ms, TR=2500 ms, flip angle=90°, 4-mm isotropic resolution, 32 axial slices). The number of collected volumes ranged from 184 to 729 across subjects (mean = 336 frames and 14.0 min).

### Image Analysis

*Preprocessing*

fMRI preprocessing in neonates included correction of intensity differences attributable to interleaved acquisition, linear realignment within and across runs to compensate for rigid body motion, bias field correction, intensity normalization of each run to a whole-brain mode value of 1000, readout distortion correction, and linear registration of BOLD images to the adult Talairach isotropic atlas (Talairach and Tournoux 1988). Volumetric data were sampled to a 3 x 3 x 3 mm space in the same step. Neonates were registered: BOLD → individual T2 → cohort-specific T2 atlas → 711-2N Talairach atlas. The cohort-specific T2 atlas was generated from a subset of 50 neonates from the eLABE dataset. Linear registrations were performed in a single step (Smith et al. 2004). Field distortion correction was performed using the FSL TOPUP toolbox (http://fsl.fmrib.ox.ac.uk/fsl/fslwiki/TOPUP). Prior to FC processing, cortical volumetric preprocessed BOLD data were mapped to subject-specific surfaces using established procedures adapted from the Human Connectome Project (Marcus et al. 2011, 2013). Using MCRIBS, each subject's surface was created from a T2-weighted anatomical image that had been linearly transformed to adult Talairach space. Therefore, the 3D coordinate systems for preprocessed volumetric BOLD data and the anatomical surface are the same. Subject-specific surfaces were nonlinearly aligned to a common surface space (fsLR_32k) using spherical registration (Glasser et al. 2013).

fMRI preprocessing in children was performed according to the ABCD-HCP BIDS fMRI pipeline (Feczko et al. 2021), which included some deviations from the original HCP pipeline (Glasser et al. 2013). fMRI preprocessing in children included correction of gradient-nonlinearity-induced distortions to the echo planar images, intensity normalization to a whole-brain mode value of 1,000, within run correction for head movement, and nonlinear registration of BOLD images to the adult Montreal Neurological Institute (MNI) space. Children were first registered: BOLD → individual T1 → adult MNI152 atlas. Children data were subsequently transformed to the 711-2B Talairach atlas and re-sampled to 3 x 3 x 3 mm space. Field distortion correction was performed using the FSL TOPUP toolbox (http://fsl.fmrib.ox.ac.uk/fsl/fslwiki/TOPUP). Prior to FC processing, cortical volumetric preprocessed BOLD data were mapped to subject-specific surfaces using established procedures adapted from the Human Connectome Project (Marcus et al. 2011, 2013). Once mapped to the surface, BOLD data were resampled to a common surface space (fsLF_32k) in a single step. These surfaces were combined with volumetric subcortical data using Connectome Workbench and smoothed with a 2mm full-width-half-maximum kernel applied to geodesic distances on surface data and Euclidean distances on volumetric data.

fMRI preprocessing in adults included correction of intensity differences attributable to interleaved acquisition, linear realignment within and across runs to compensate for rigid body motion, intensity normalization of each run to a whole-brain mode value of 1,000, and linear registration of BOLD images to the adult Talairach isotropic atlas (Talairach and Tournoux 1988). Volumetric data were sampled to a 3 x 3 x 3 mm space in the same step. Adults were registered: BOLD → individual T2 →

individual T1 → 711-2B Talairach atlas. Linear registrations were performed in a single step (Smith et al. 2004). Prior to FC processing, cortical volumetric preprocessed BOLD data were mapped to subject-specific surfaces using established procedures adapted from the Human Connectome Project (Marcus et al. 2011, 2013). Using Freesurfer, each subject's surface was created from a T1-weighted anatomical image that had been linearly transformed to adult Talairach space. Subject-specific surfaces were nonlinearly aligned to a common surface space (fsLR_32k) using spherical registration (Glasser et al. 2013).

*FC Processing*

In the neonates, pre-processed cortical data were first mapped to the surface (see above), smoothed with a small geodesic 2D Gaussian kernel ($\sigma$=1mm), and then processed in surface-space. Child cortical data were also processed in surface-space. In the adults, FC processing was done in volume-space, although FC processed data were mapped to subject-specific surfaces prior to further FC analyses and subject averaging, so the adult analyses are surface-aligned. Temporal masks were created that censored high-motion frames based on study-specific protocols (see below). Censored frames were ignored in FC processing, which included the following steps: (i) demean and detrend within run, (ii) multiple regression with nuisances timeseries including white matter, ventricles, and whole brain (neonates: average gray matter signal, children and adults: whole brain mask), as well as 24 parameters derived from head motion. Finally, retained data were interpolated into censored timepoints to allow band-pass filtering (0.005 Hz < f < 0.1 Hz for neonates; 0.008 Hz < f < 0.09 using a 2$^{nd}$ order Butterworth filter for children; 0.009 Hz < f < 0.08 Hz for adults). After FC processing, time courses for neonatal and adult volumetric subcortical data were combined with surface data and smoothed with geodesic 2D Gaussian and Euclidean 3D kernels ($\sigma$=2.25mm for neonates and $\sigma$=2.55mm for adults).

*Motion Scrubbing*

As recommended, FD thresholds were tailored for each dataset to account for differences in head size, respiratory rate, and temporal sampling rate (i.e., TR; Power et al. 2012, 2013, 2014). Prior to motion censoring in the children, the FD trace was filtered to remove respiratory signal (18.562 to 25.726 breaths per minute). Volumes were included only in temporally contiguous sets of at least 3 volumes for neonates and 5 volumes for adults. BOLD runs were included only if they retained a minimum of 130 such frames for neonates and 30 such frames for adults. There was no minimum number of contiguous volumes or usable frames per BOLD run for the children.

**Anatomical Definitions**

*Subcortical Definitions*

Basal ganglia and thalamus segmentations for neonates were derived from an atlas generated from an average T2 of 50 subjects. The neonatal atlas was segmented using MCRIBS and then manually edited. Subcortical masks were then applied to individuals. For children and adults, a template segmentation from Freesurfer was applied to atlas aligned data. For all datasets, the caudate, putamen, and pallidum were grouped together and isolated to create a basal ganglia mask. Basal ganglia and thalamus masks were then applied to individuals.

*Cortical System Definition*

Because precursors of adult functional networks are not fully mature in the neonate, we also examined the representations of baby-defined Myers-Labonte networks in the subcortex (**Supp. Fig. 4**; Myers et al. 2024). These baby-defined networks were generated using a parallel process as the Gordon networks. Additionally, to better understand the effector specificity of the somatomotor representation (e.g., face, hand, foot), we also used the Midnight Scan Club network template, which includes a somatomotor leg network as well (**Supp. Fig. 5**; Gordon et al. 2017)

**Subcortico-cortical FC Matrix Construction**

*Excluding cortical signal adjacent to the subcortex*

Anatomically, some subcortical structures are immediately adjacent to cortical regions within the 10 selected cortical networks. To account for the possibility that signal bleed between adjacent regions might lead to artifactual increases in correlation strength between subcortical voxels and certain cortical networks, we excluded vertices belonging to the 10 cortical networks that were located within 10 mm of the basal ganglia or thalamus masks from the network signal in adult Talairach space.

**Winner-Take-All Approach**

*Winner-take-all reliability*

To assess the reliability of the winner-take-all approach in our group-averaged data, we performed a bootstrapping analysis to generate 1000 resamples of each original dataset, with each resample being the same size as the original datasets. Sampling was done with replacement, such that each subject's data was returned to the dataset after being selected, allowing it to be chosen multiple times in the same resample. We first calculated the most common winner network assignment (the “mode network”) within each voxel across the 1000 iterations. We defined winner-take-all reliability for each voxel as the percentage of iterations in which that voxel was assigned the mode network. We also assessed the reliability of sensorimotor vs. association network assignments in the same way, after first designating each network as sensorimotor (visual, auditory, somatomotor hand, somatomotor face) or associative (salience, cingulo-opercular, default mode, frontoparietal, dorsal attention, ventral attention). Thus, a voxel with high sensorimotor reliability would be consistently assigned any of the sensorimotor networks across iterations.

*Cluster Stability Analysis*

We assessed the stability of clusters for k=2 through k=9 in the neonatal dataset to determine a reasonable k value for the basal ganglia and thalamus (Lange et al. 2004; Sylvester et al. 2020). For each k value, we performed 1000 times iterations in which the total number of basal ganglia or thalamus voxels were randomly divided into two equal split-halves (subset A and subset B). K-means clustering was performed separately on each half. Then, the voxels from subset B were additionally assigned to the classification derived from the k-means clustering of subset A by assigning each voxel in subset B to the closest cluster center in subset A. Cluster center refers to the average vertexwise connectivity to cortex of the cluster. Next, we computed the Hamming distance between the vector of the assignments of subset B based on the closest center from subset A. The Hamming distance was computed for all possible permutations of labels of the solution derived from the k-means clustering of subset B, since the labels are arbitrary. The raw stability value for each iteration was defined as the lowest Hamming distance computed from these permutations. Raw stability values were normalized by a 'randomized stability value' that was calculated separately for each k value to account for the fact that the chance misclassification rate varies as a function of k. This randomized stability value was calculated by randomly assigning all voxels in subset B to k clusters two separate times and then deriving the Hamming distance of these randomized assignments (including permuting labels and performing 1000 iterations as above). Finally, we calculated and graphed the slope/derivative of the normalized stability values to determine how increasing the number of clusters changes the stability of the groupings. We concluded that k=4 was optimal for both the basal ganglia and thalamus to minimize the number of clusters while maximizing the change in stability from increasing to a greater cluster number.

### SUPPLEMENTARY FIGURES

| | BASAL GANGLIA | | | THALAMUS | | |
|---|---|---|---|---|---|---|
| | Neonates v. Children | Neonates v. Adults | Children v. Adults | Neonates v. Children | Neonates v. Adults | Children v. Adults |
| **VIS** | N/A | N/A | n.s. | p<0.001 | p<0.001 | p<0.05 |
| **AUD** | N/A | N/A | n.s. | n.s. | p<0.001 | p<0.001 |
| **SMH** | n.s. | p<0.001 | p<0.05 | n.s. | p<0.001 | p<0.001 |
| **SMF** | n.s. | n.s. | n.s. | n.s. | p<0.001 | p<0.001 |
| **SAL** | p<0.001 | p<0.001 | p<0.001 | p<0.001 | p<0.001 | p<0.001 |
| **CON** | p<0.001 | p<0.001 | p<0.001 | n.s. | p<0.001 | p<0.001 |
| **FPN** | p<0.001 | p<0.001 | p<0.001 | n.s. | p<0.001 | p<0.001 |
| **DMN** | p<0.001 | p<0.01 | p<0.05 | p<0.01 | p<0.001 | p<0.05 |
| **DAN** | n.s. | n.s. | n.s. | n.s. | p<0.001 | p<0.001 |
| **VAN** | N/A | N/A | N/A | N/A | N/A | N/A |

**Supplementary Table 1.** *Statistical significance of age-related differences in network selectivity for the basal ganglia (left) and the thalamus (right)*. Network selectivity was calculated as the difference between the FC strength of the winner network and the average FC strength of all other networks within each network-selective subcortical ROI. Mean winner network selectivity was compared across age groups with two-sample t-tests. False discovery rate was used to correct for the multiple comparisons (10 networks). A value of "N/A" indicates that one of the groups had no network-selective subcortical ROI for the given network. A value of "n.s." indicates no significant difference between the age groups. Directionality of the effect: red indicates selectivity is greater in the older age group; blue indicates selectivity is greater in the younger age group. See also **Figure 4** in the main text.

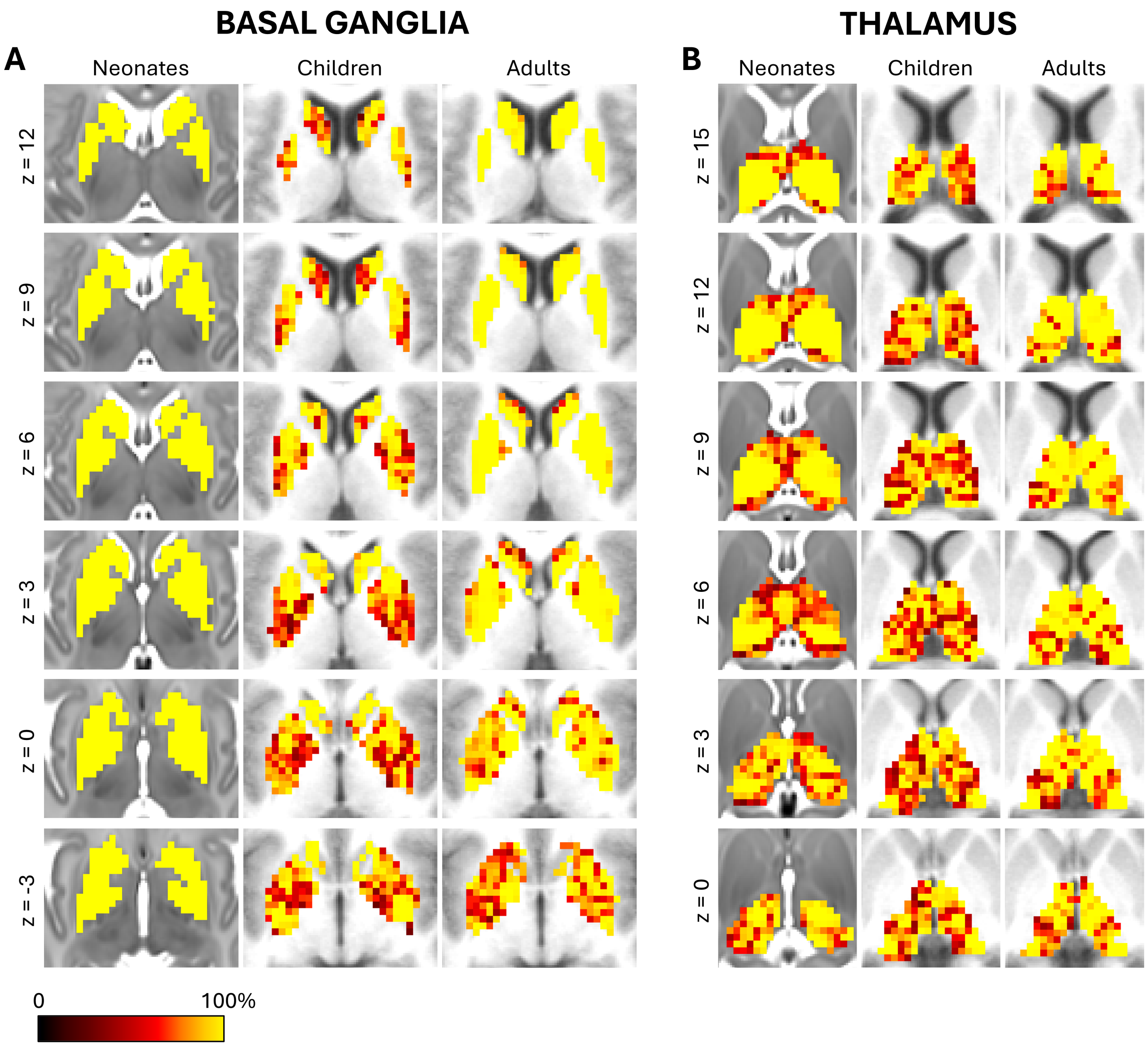


**Supplementary Figure 1***. Reliability of group-averaged winner assignments in **A)** the basal ganglia and **B)** the thalamus in neonates, children, and adults.* Reliability was calculated as the percentage of bootstrap iterations in which each voxel was assigned the mode winner network. Slices are shown in the transverse plane (image left is anatomical left), z = -3 to z = 12 for the basal ganglia and z = 0 to z = 15 for the thalamus. Group-averaged winner-take-all maps are highly reliable with the given number of subjects within each age group (average reliability – neonatal BG: 100%, neonatal THAL: 86.51%, child BG: 80.56%, child THAL: 79.20%, adult BG: 90.07%, adult THAL: 88.20%).

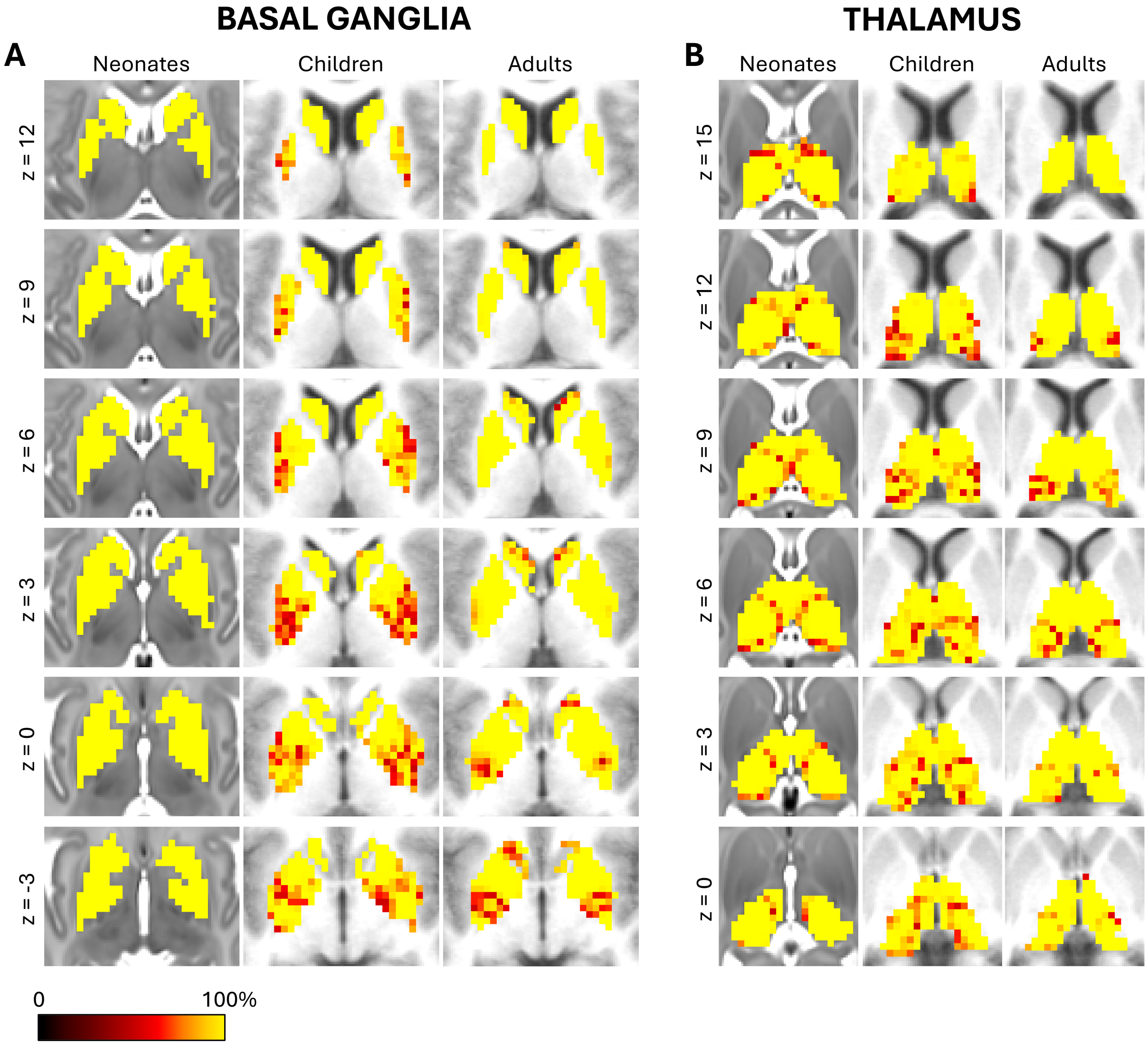


**Supplementary Figure 2***. Sensorimotor vs. association reliability of group-averaged winner assignments in* ***A)*** *the basal ganglia and* ***B)*** *the thalamus in neonates, children, and adults.* Sensorimotor vs. association reliability was defined as how in agreement each voxel was across bootstrap iterations for sensorimotor or association network preference. Slices are shown in the transverse plane (image left is anatomical left), z = -3 to z = 12 for the basal ganglia and z = 0 to z = 15 for the thalamus. Average sensorimotor vs. association reliability – neonatal BG: 100%, neonatal THAL: 95.75%, child BG: 92.74%, child THAL: 93.53%, adult BG: 95.77%, adult THAL: 96.79%.

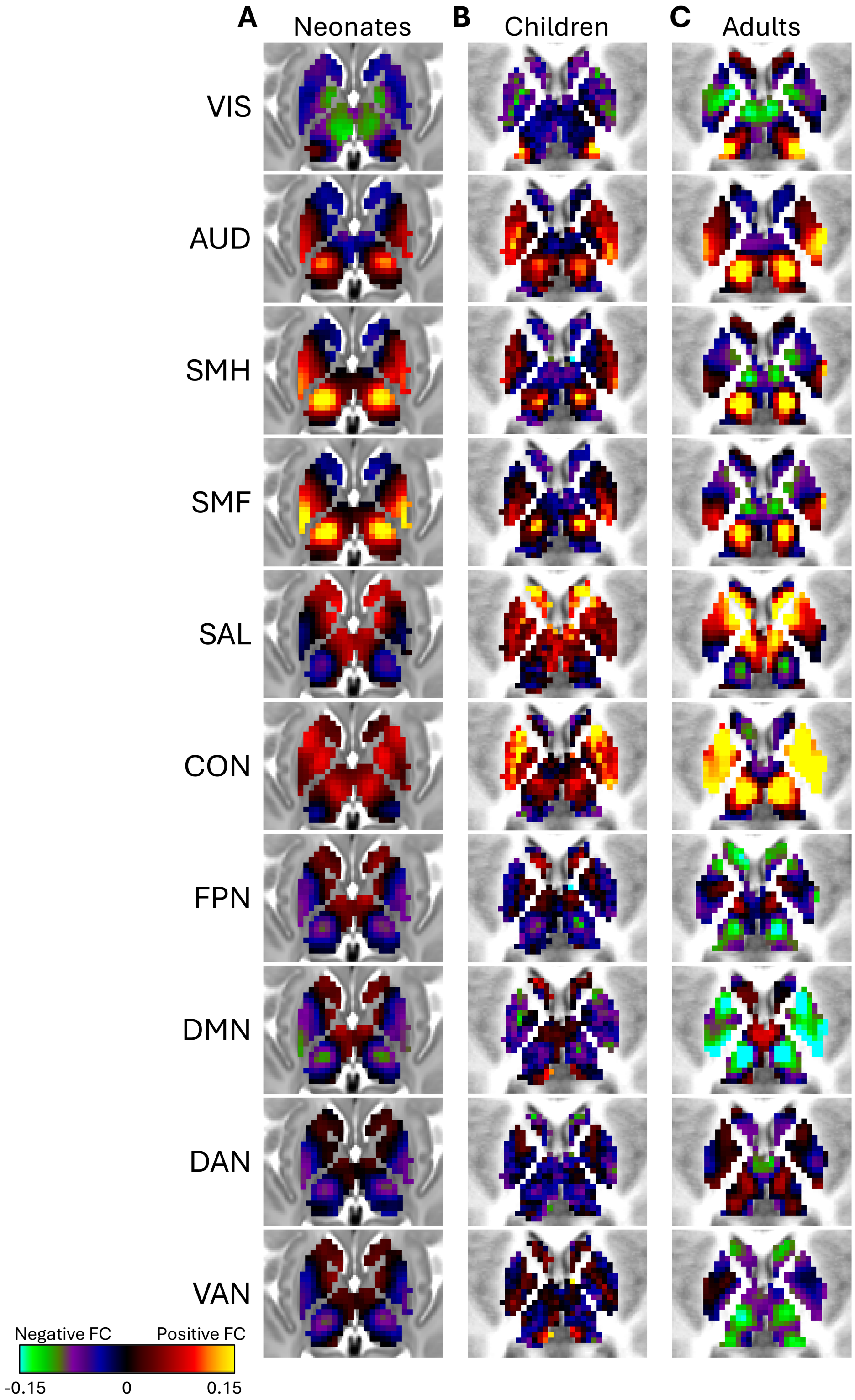


**Supplementary Figure 3***. Subcortical FC with each adult cortical network for* ***A)*** *neonates,* ***B)*** *children, and* ***C)*** *adults.* All slices are shown in the transverse plane at z = 3. Image left is anatomical left. Network labels: VIS, AUD, SMH, SMF, SAL, CON, FPN, DMN, DAN, VAN.

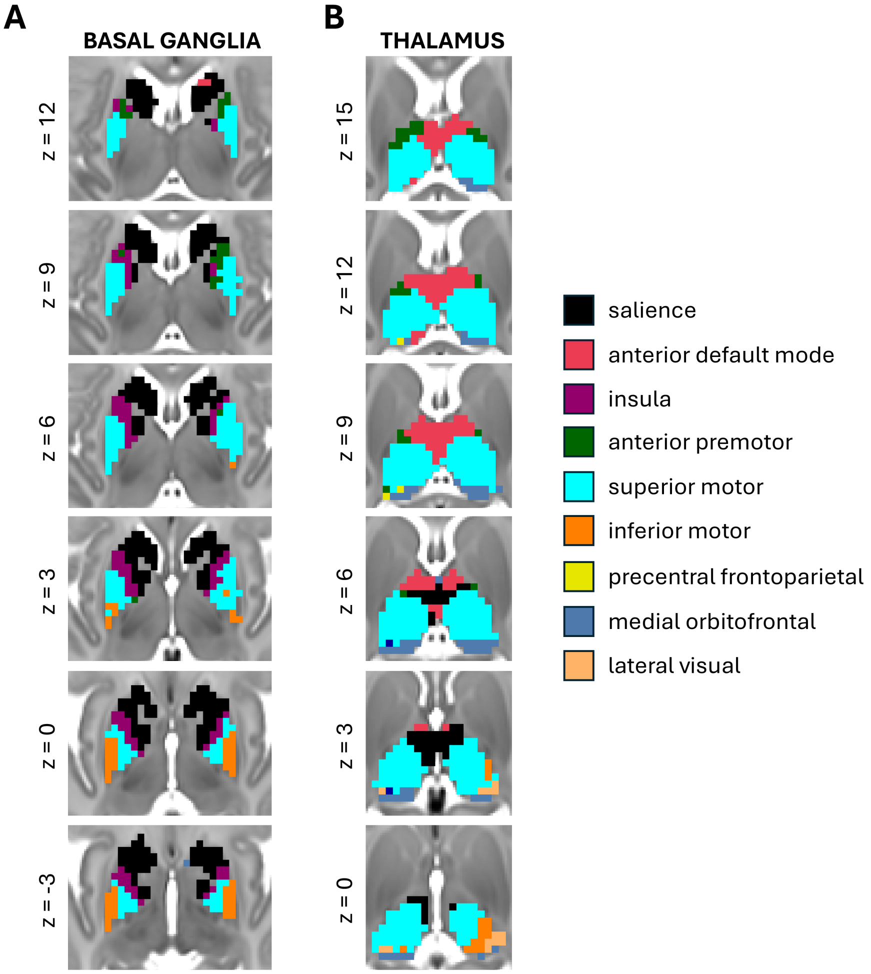


**Supplementary Figure 4***. Functional network organization in the neonatal* ***A)*** *basal ganglia and* ***B)*** *thalamus based on the Myers-Labonte networks* (Myers et al. 2024)*.* The Myers-Labonte network scheme divides many association networks into their anterior and posterior components. Each subcortical voxel is colored according to the cortical network with which it has the highest FC (color key on the right). These maps demonstrate that the neonatal basal ganglia and thalamus exhibit a preference for anterior networks. Slices are shown in the transverse plane (image left is anatomical left), z = -3 to z = 12 for the basal ganglia and z = 0 to z = 15 for the thalamus.

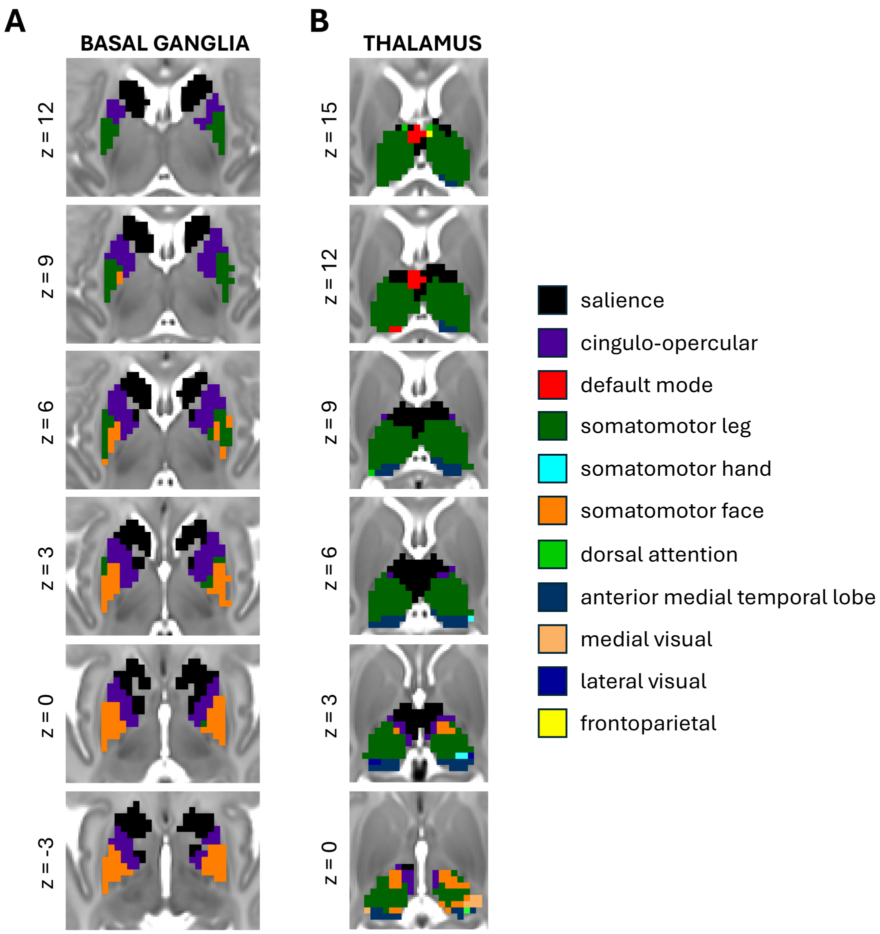


**Supplementary Figure 5***. Functional network organization in the neonatal* ***A)*** *basal ganglia and* ***B)*** *thalamus based on the Midnight Scan Club networks* (Gordon et al. 2017). The Midnight Scan Club network template differentiates between the somatomotor hand and leg effector networks. Each subcortical voxel is colored according to the cortical network with which it has the highest FC (color key on the right). These maps demonstrate that the neonatal basal ganglia and thalamus exhibit preferential connectivity to the somatomotor leg network compared to the somatomotor hand network. Slices are shown in the transverse plane (image left is anatomical left), z = -3 to z = 12 for the basal ganglia and z = 0 to z = 15 for the thalamus.

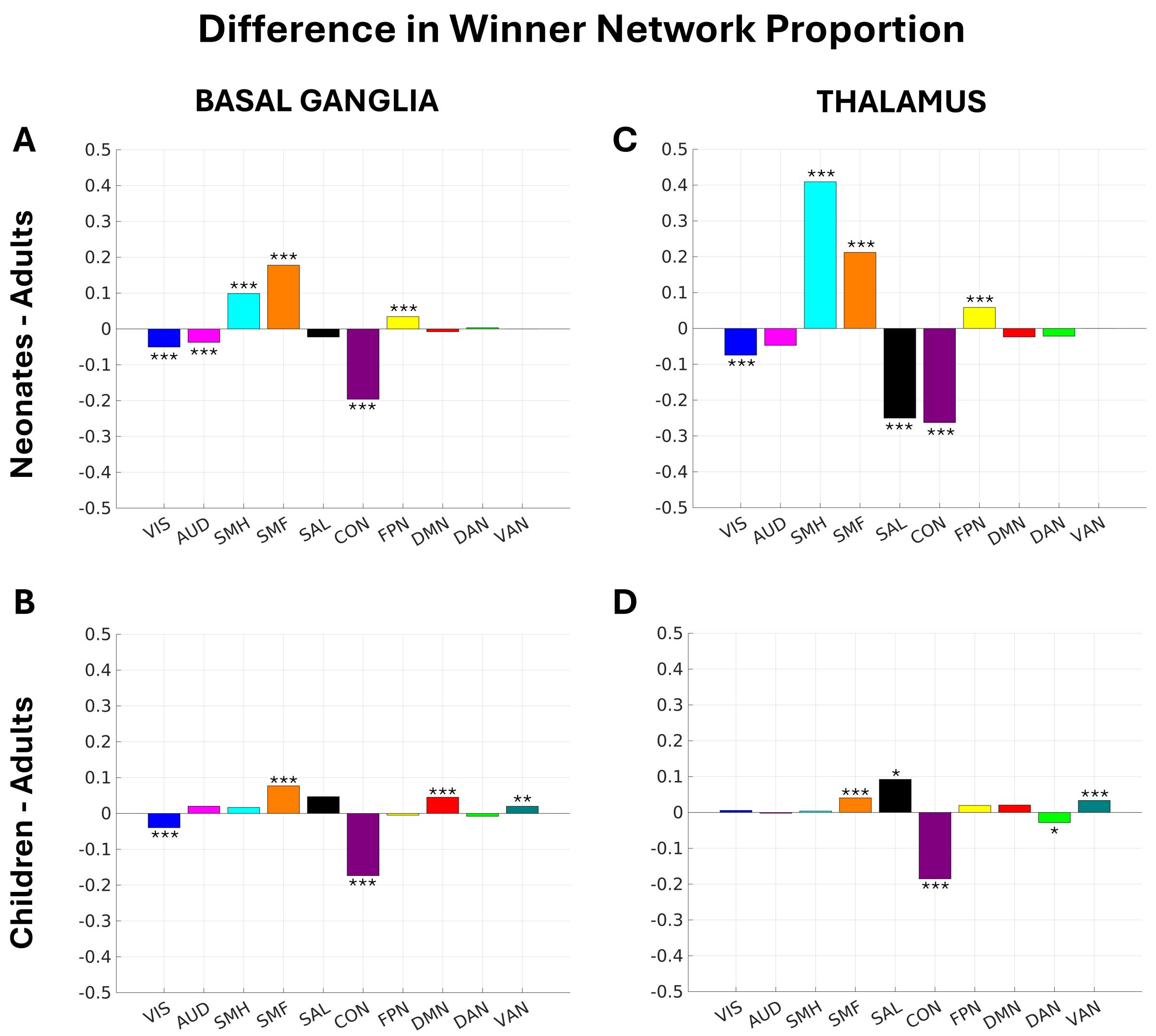


**Supplementary Figure 6***. Bar graphs displaying age-related differences in the relative sizes of network-selective ROIs in* ***A, B)*** *the basal ganglia and* ***C, D)*** *the thalamus*. **A, C)** Bar graphs comparing neonates and adults. A bar above the x-axis indicates that the network-selective ROI is larger in neonates than in adults. A bar below the x-axis indicates that the network-selective ROI is smaller in neonates than in adults. **B, D)** Bar graphs comparing children and adults. A bar above the x-axis indicates that the network-selective ROI is larger in children than in adults. A bar below the x-axis indicates that the network-selective ROI is smaller in children than in adults. Significant differences in size were determined via bootstrapping. *p<0.05, **p<0.01, ***p<0.001.

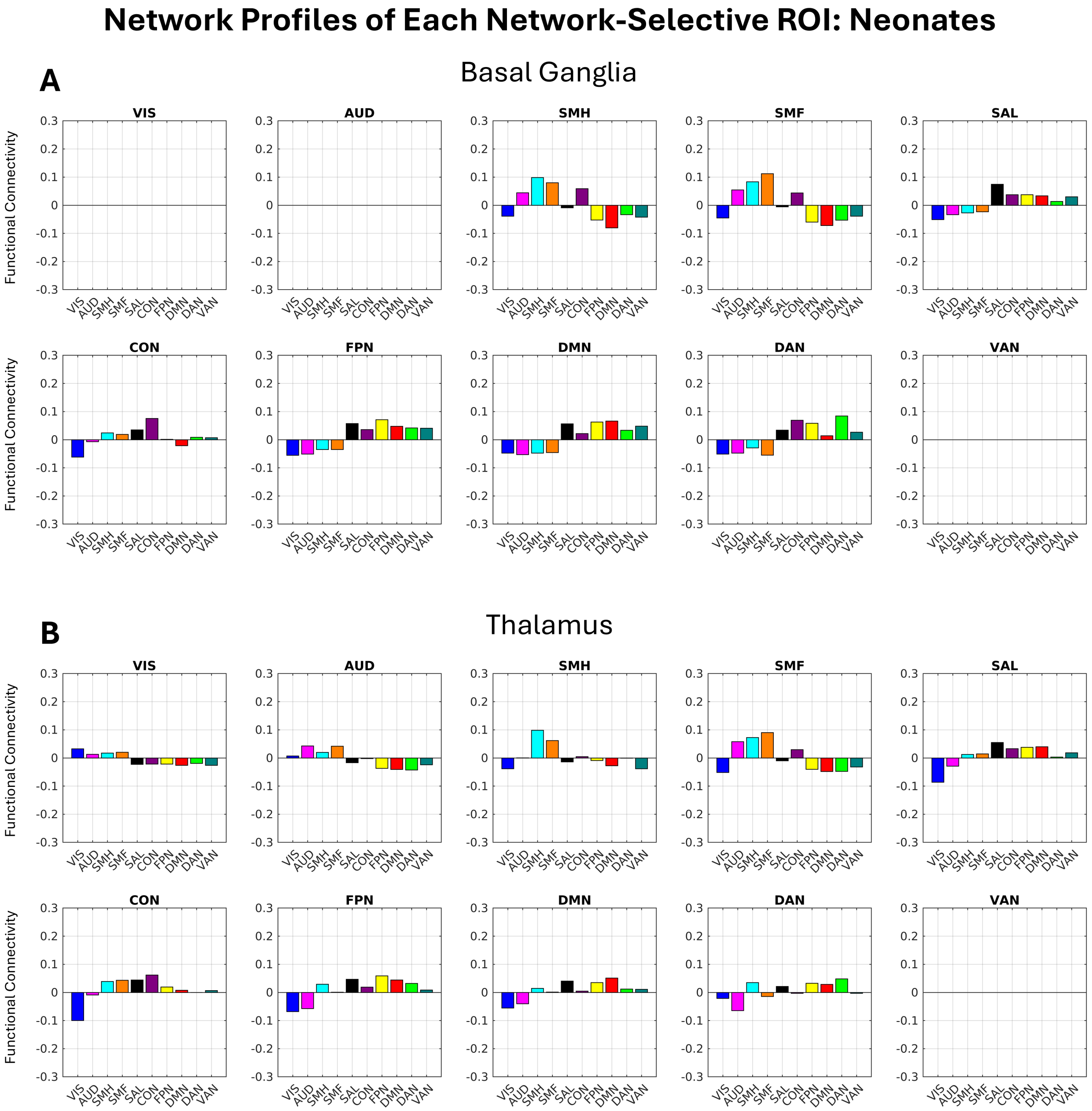


**Supplementary Figure 7***. Bar graphs displaying FC to each network for each network-selective ROI in the* ***A)*** *basal ganglia and* ***B)*** *thalamus in neonates.* Each subplot represents the average FC to each of the 10 Gordon networks across the voxels within the given network's network-selective subcortical ROI. Each network exhibits the strongest FC compared to other networks within its own network-selective subcortical ROI.

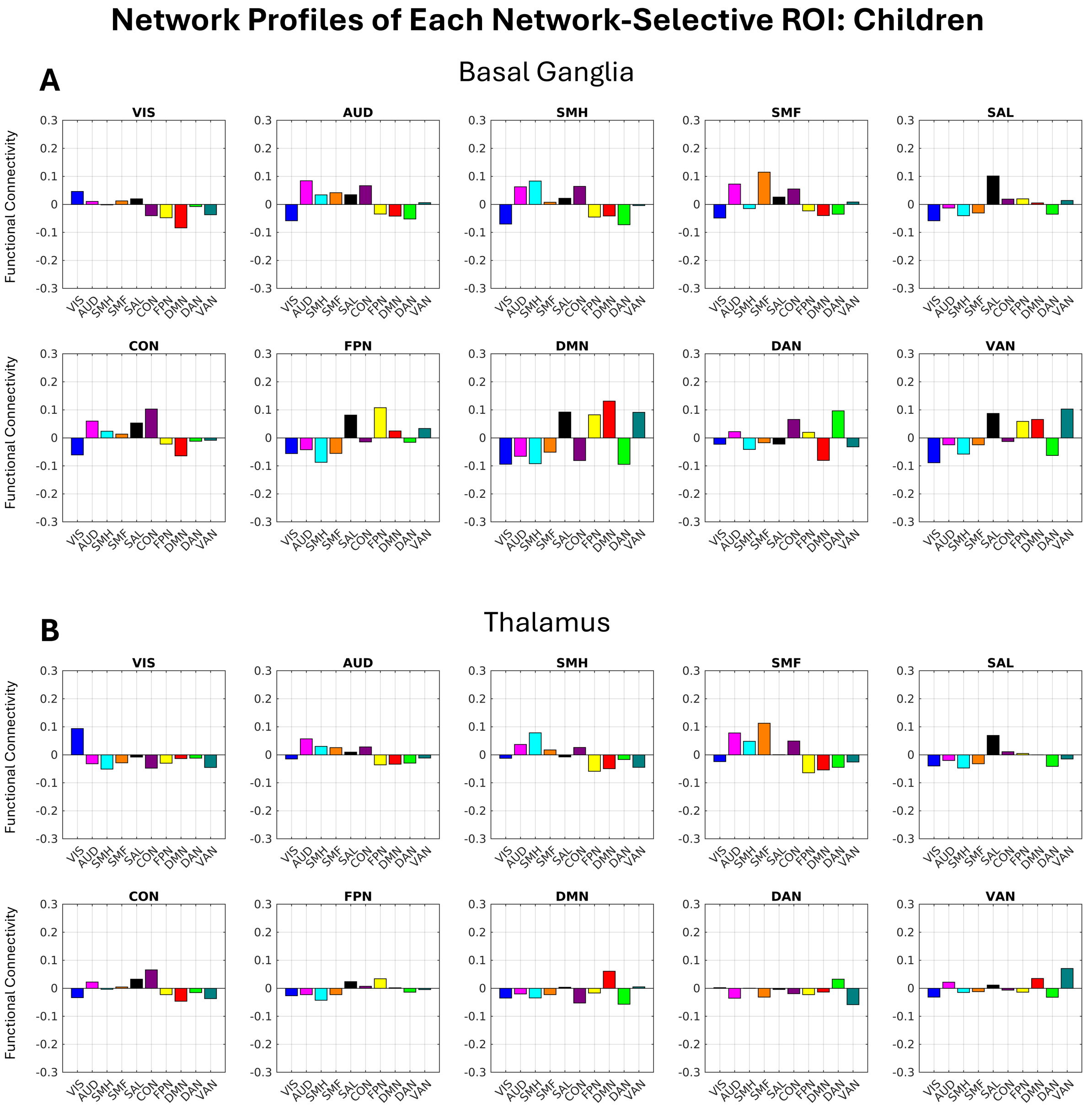


**Supplementary Figure 8***. Bar graphs displaying FC to each network for each network-selective ROI in the* ***A)*** *basal ganglia and* ***B)*** *thalamus in children.* Each subplot represents the average FC to each of the 10 Gordon networks across the voxels within the given network's network-selective subcortical ROI. Each network exhibits the strongest FC compared to other networks within its own network-selective subcortical ROI.

### Network Profiles of Each Network-Selective ROI: Adults

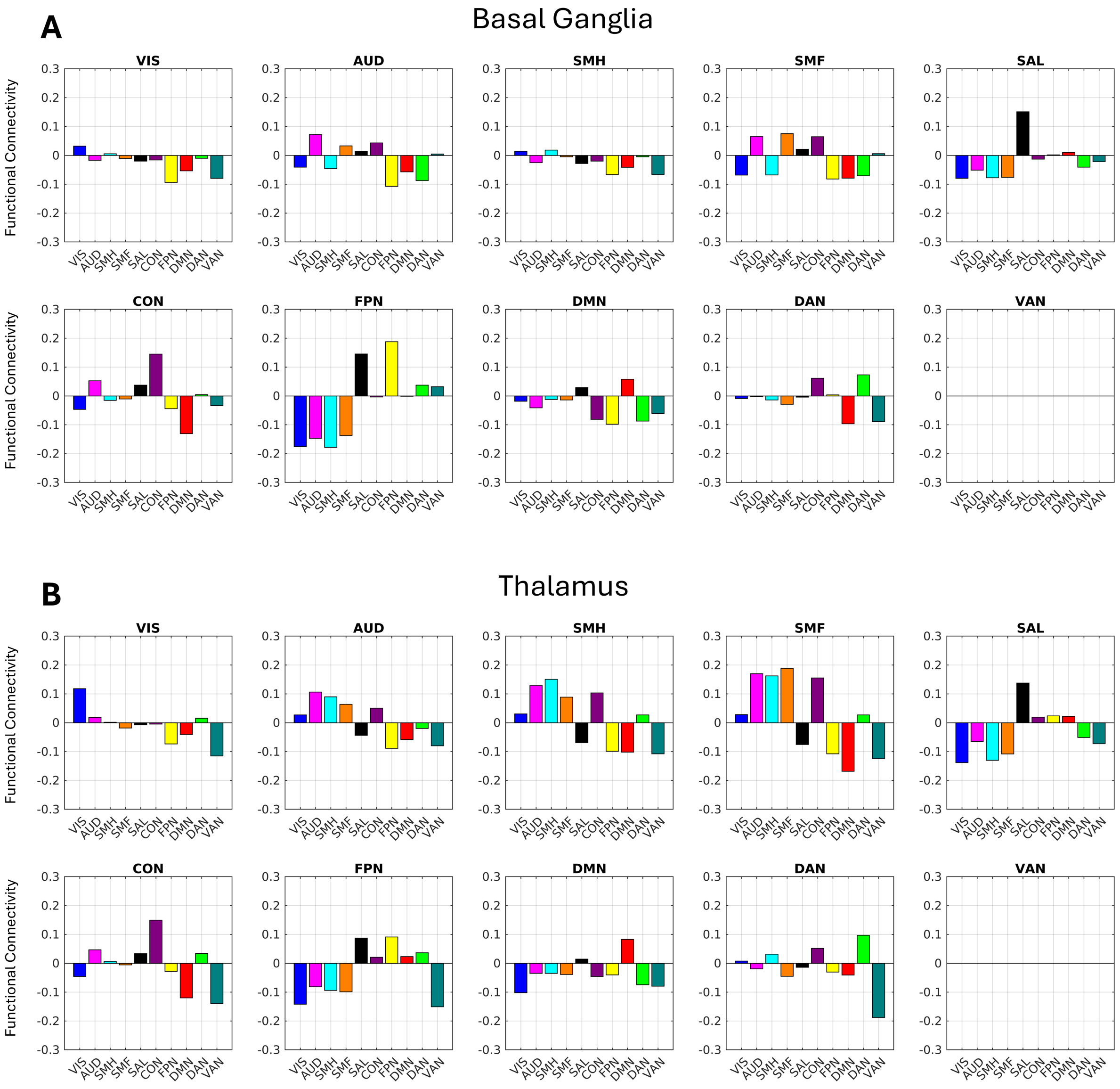


**Supplementary Figure 9***. Bar graphs displaying FC to each network for each network-selective ROI in the* ***A)*** *basal ganglia and* ***B)*** *thalamus in adults.* Each subplot represents the average FC to each of the 10 Gordon networks across the voxels within the given network's network-selective subcortical ROI. Each network exhibits the strongest FC compared to other networks within its own network-selective subcortical ROI.

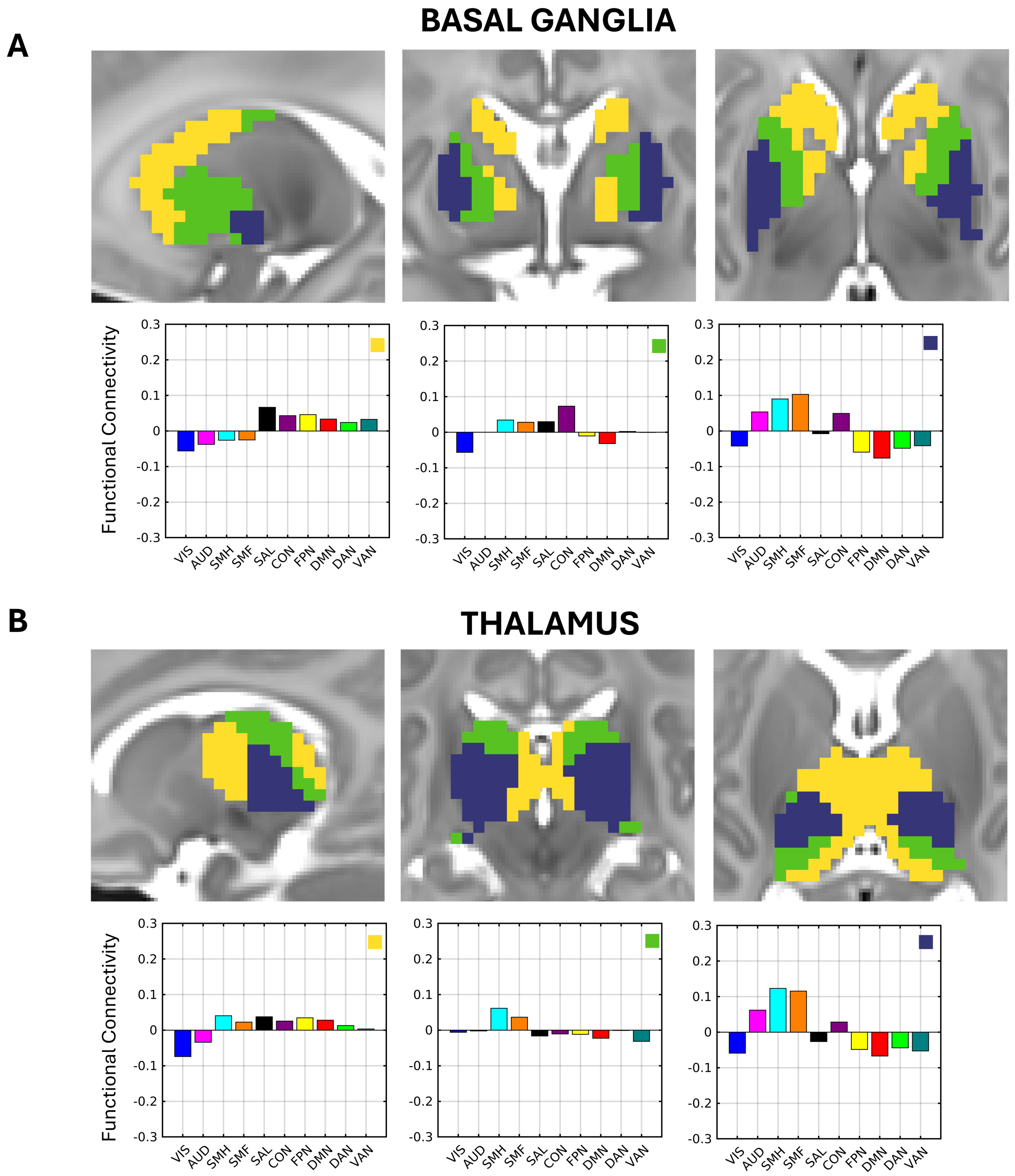


**Supplementary Figure 10***. K=3 clustering results for the neonatal* ***A****) basal ganglia and* ***B)*** *thalamus according to FC profiles with the Gordon networks.* Sagittal (left), coronal (middle), and transverse (right) slices are displayed. Basal ganglia coordinates: x = -20, y = -3, z = 3. Thalamus coordinates: x = -12, y = -24, z = 9. Average network profiles showing average FC to each network for the voxels within each cluster are depicted below. Each network profile graph contains a box with its corresponding cluster color in the top right.

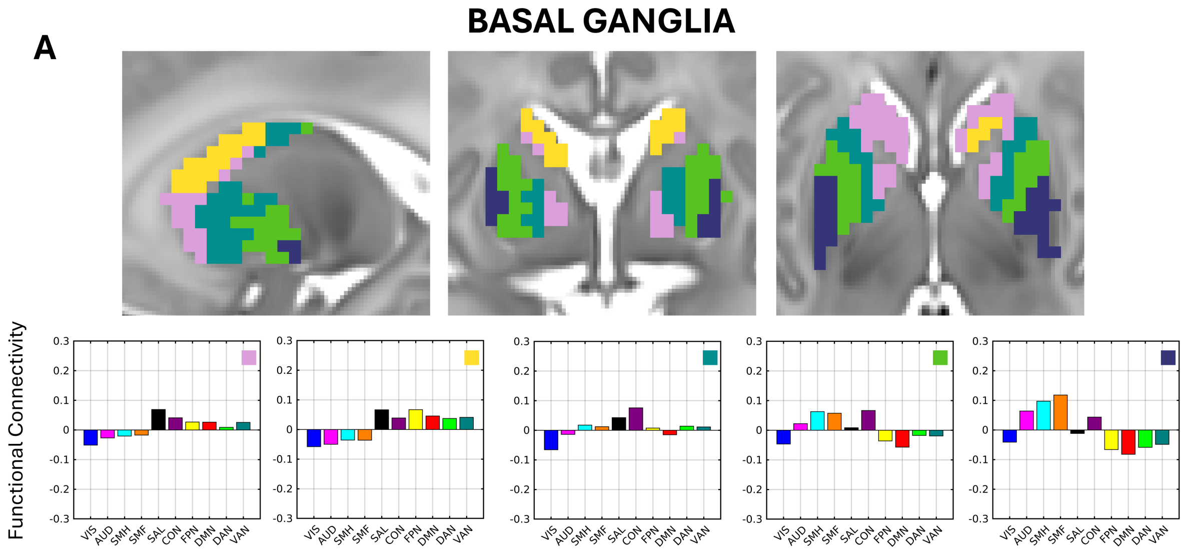


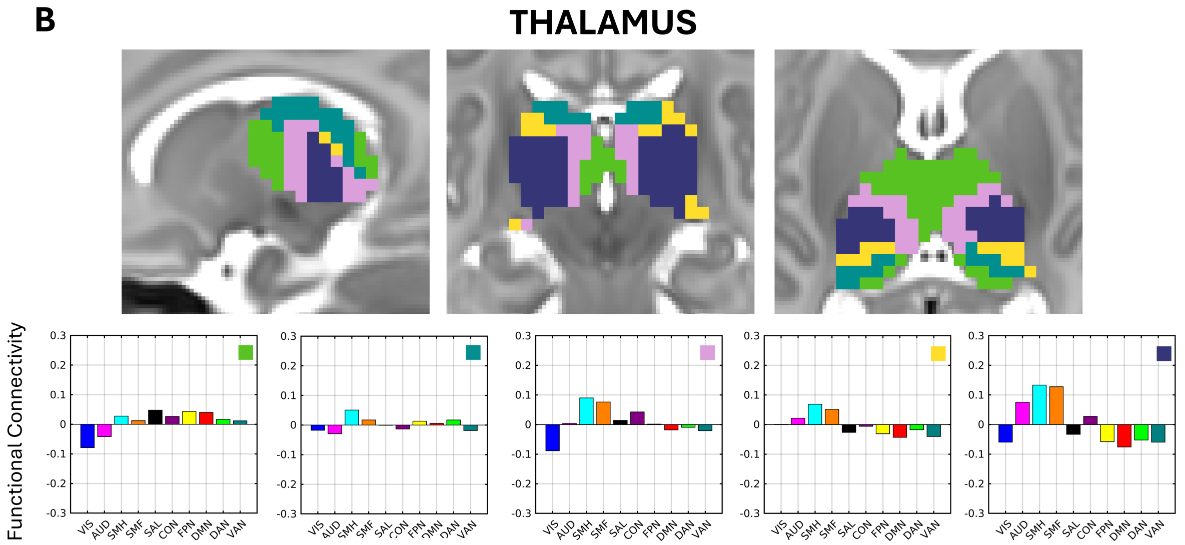


**Supplementary Figure 11***. K=5 clustering results for the neonatal* ***A****) basal ganglia and* ***B)*** *thalamus according to FC profiles with the Gordon networks.* Sagittal (left), coronal (middle), and transverse (right) slices are displayed. Basal ganglia coordinates: x = -20, y = -3, z = 3. Thalamus coordinates: x = -12, y = -24, z = 9. Average network profiles showing average FC to each network for the voxels within each cluster are depicted below. Each network profile graph contains a box with its corresponding cluster color in the top right.

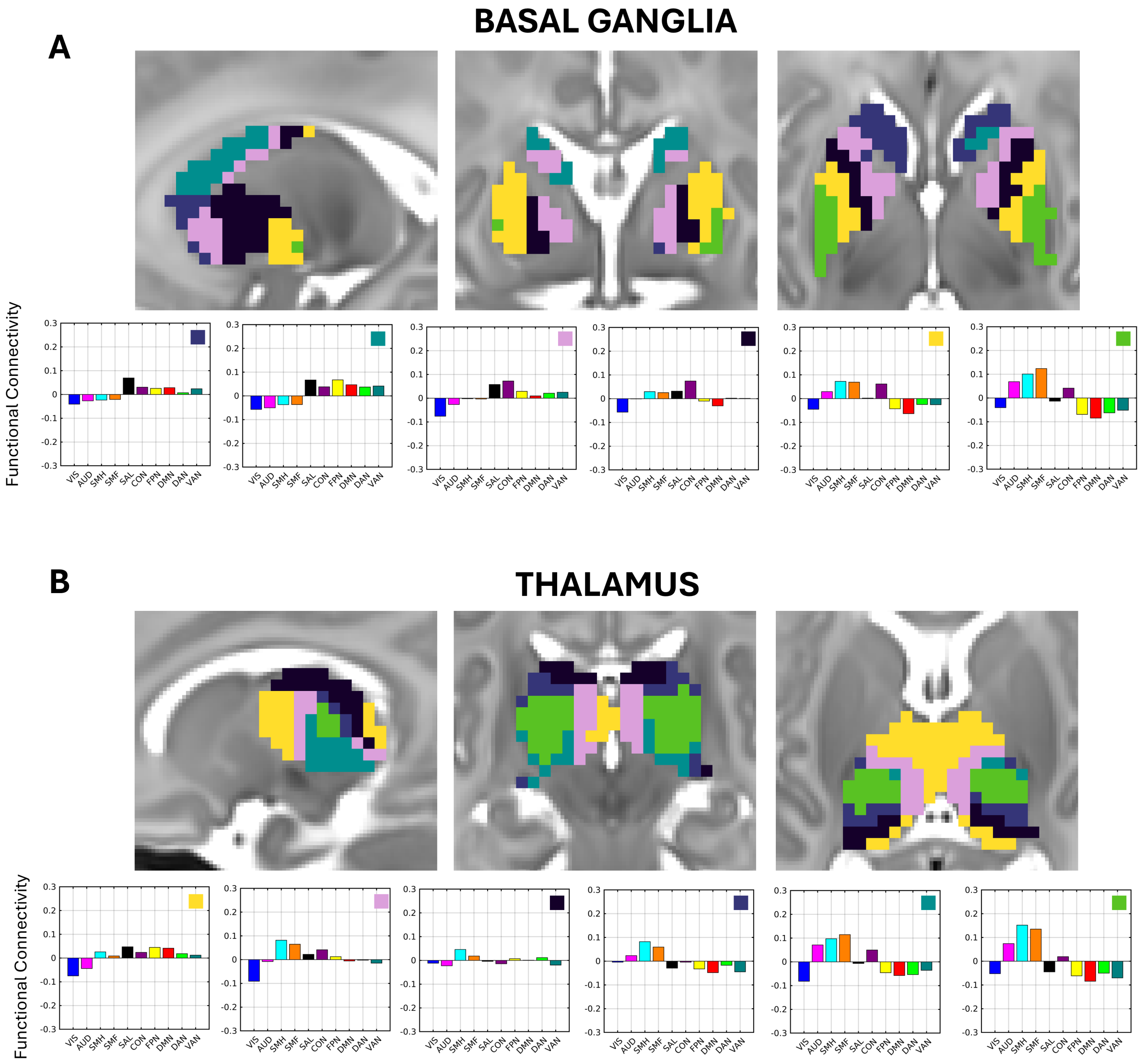


**Supplementary Figure 12***. K=6 clustering results for the neonatal* ***A****) basal ganglia and* ***B)*** *thalamus according to FC profiles with the Gordon networks.* Sagittal (left), coronal (middle), and transverse (right) slices are displayed. Basal ganglia coordinates: x = -20, y = -3, z = 3. Thalamus coordinates: x = -12, y = -24, z = 9. Average network profiles showing average FC to each network for the voxels within each cluster are depicted below. Each network profile graph contains a box with its corresponding cluster color in the top right.

**A**

### BASAL GANGLIA

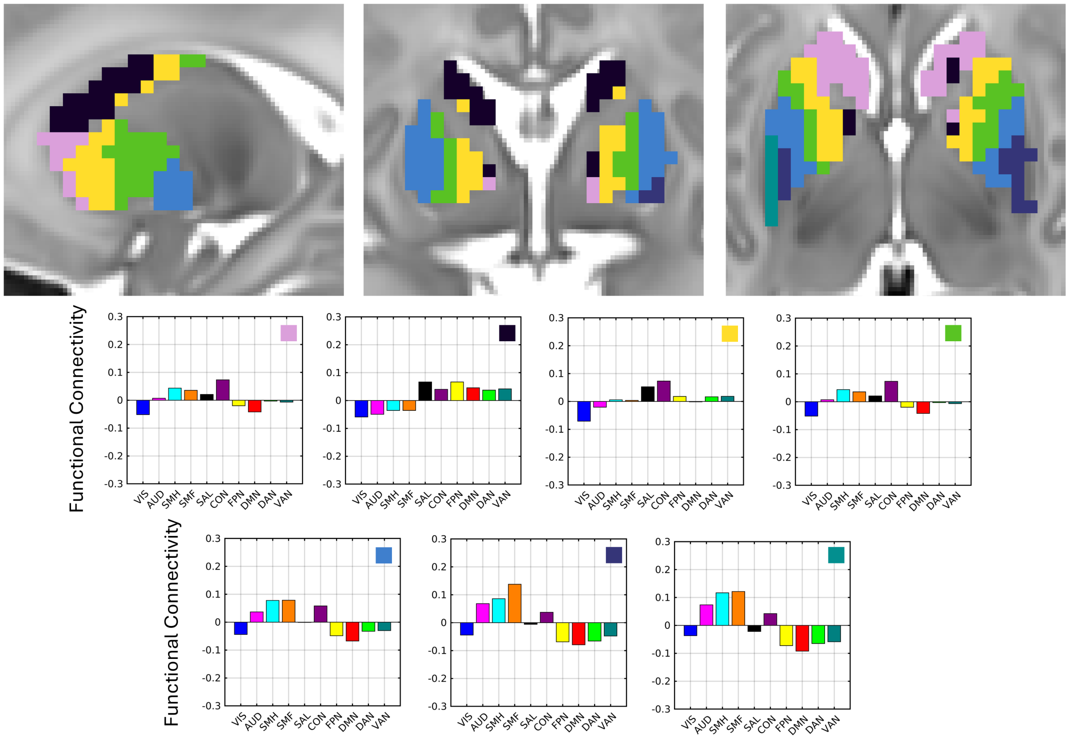


**B**

### THALAMUS

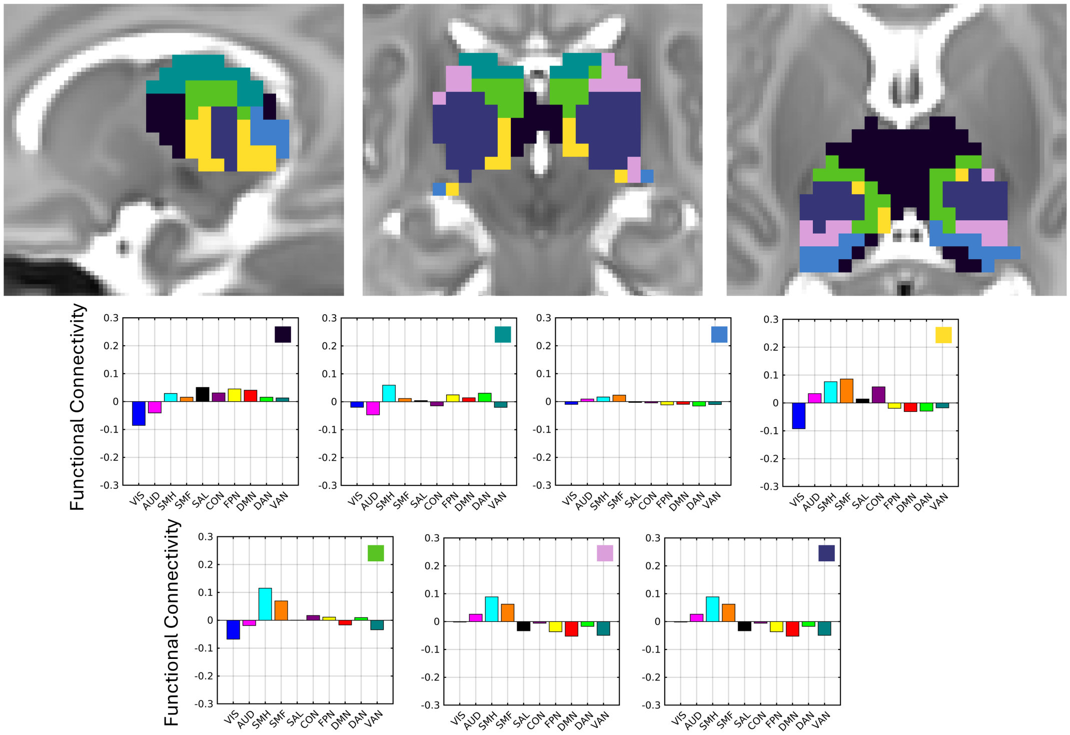

**Supplementary Figure 13***. K=7 clustering results for the neonatal* ***A****) basal ganglia and* ***B)*** *thalamus according to FC profiles with the Gordon networks.* Sagittal (left), coronal (middle), and transverse (right) slices are displayed. Basal ganglia coordinates: x = -20, y = -3, z = 3. Thalamus coordinates: x = -12, y = -24, z = 9. Average network profiles showing average FC to each network for the voxels within each cluster are depicted below. Each network profile graph contains a box with its corresponding cluster color in the top right.

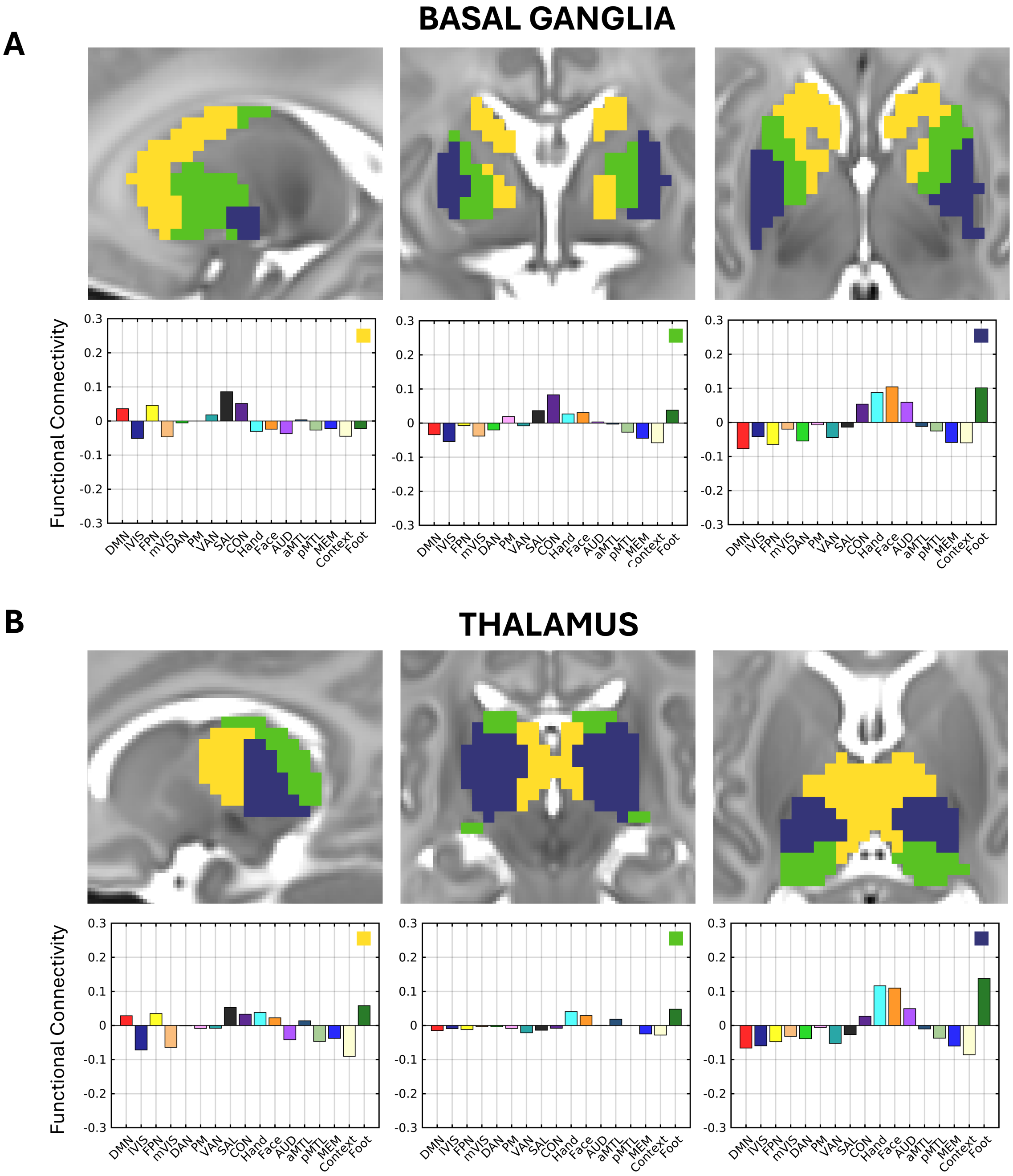


**Supplementary Figure 14.** *K=3 clustering results for the neonatal **A**) basal ganglia and **B)** thalamus according to FC profiles with the Midnight Scan Club networks.* Sagittal (left), coronal (middle), and transverse (right) slices are displayed. Basal ganglia coordinates: x = -20, y = -3, z = 3. Thalamus coordinates: x = -12, y = -24, z = 9. Average network profiles showing average FC to each network for the voxels within each cluster are depicted below. Each network profile graph contains a box with its corresponding cluster color in the top right.

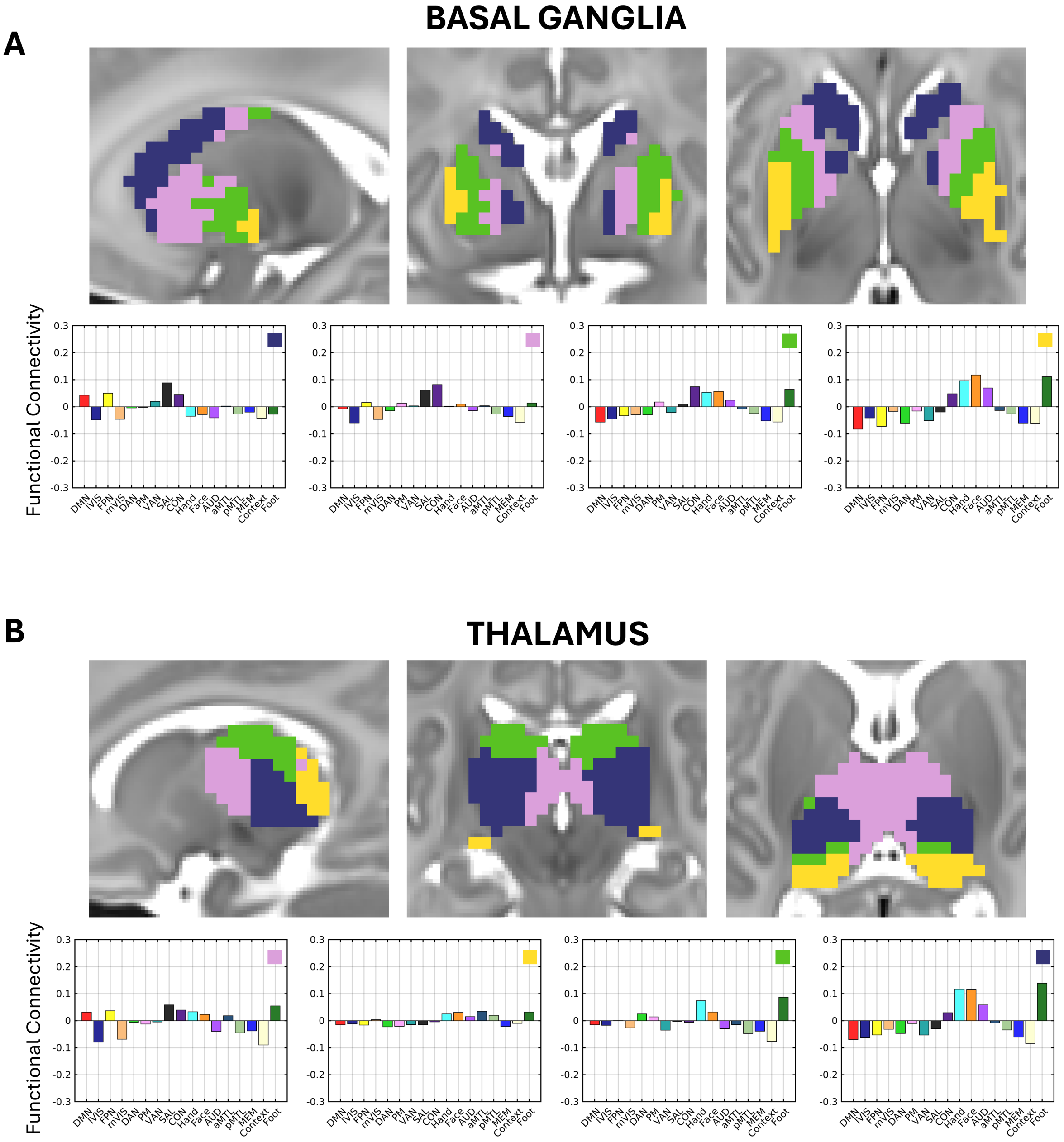


**Supplementary Figure 15.** *K=4 clustering results for the neonatal **A**) basal ganglia and **B)** thalamus according to FC profiles with the Midnight Scan Club networks.* Sagittal (left), coronal (middle), and transverse (right) slices are displayed. Basal ganglia coordinates: x = -20, y = -3, z = 3. Thalamus coordinates: x = -12, y = -24, z = 9. Average network profiles showing average FC to each network for the voxels within each cluster are depicted below. Each network profile graph contains a box with its corresponding cluster color in the top right.

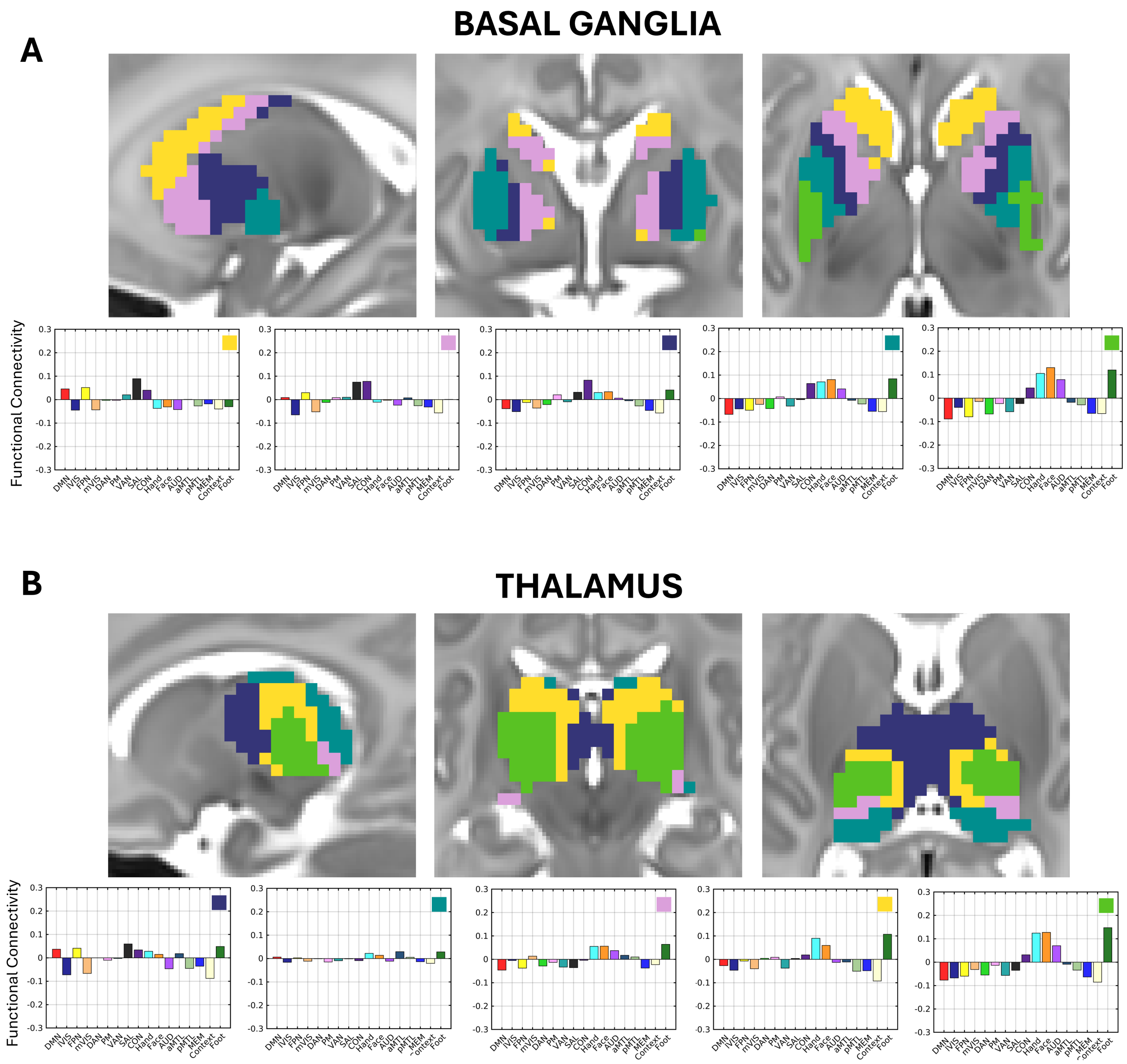


**Supplementary Figure 16.** *K=5 clustering results for the neonatal **A**) basal ganglia and **B)** thalamus according to FC profiles with the Midnight Scan Club networks.* Sagittal (left), coronal (middle), and transverse (right) slices are displayed. Basal ganglia coordinates: x = -20, y = -3, z = 3. Thalamus coordinates: x = -12, y = -24, z = 9. Average network profiles showing average FC to each network for the voxels within each cluster are depicted below. Each network profile graph contains a box with its corresponding cluster color in the top right.

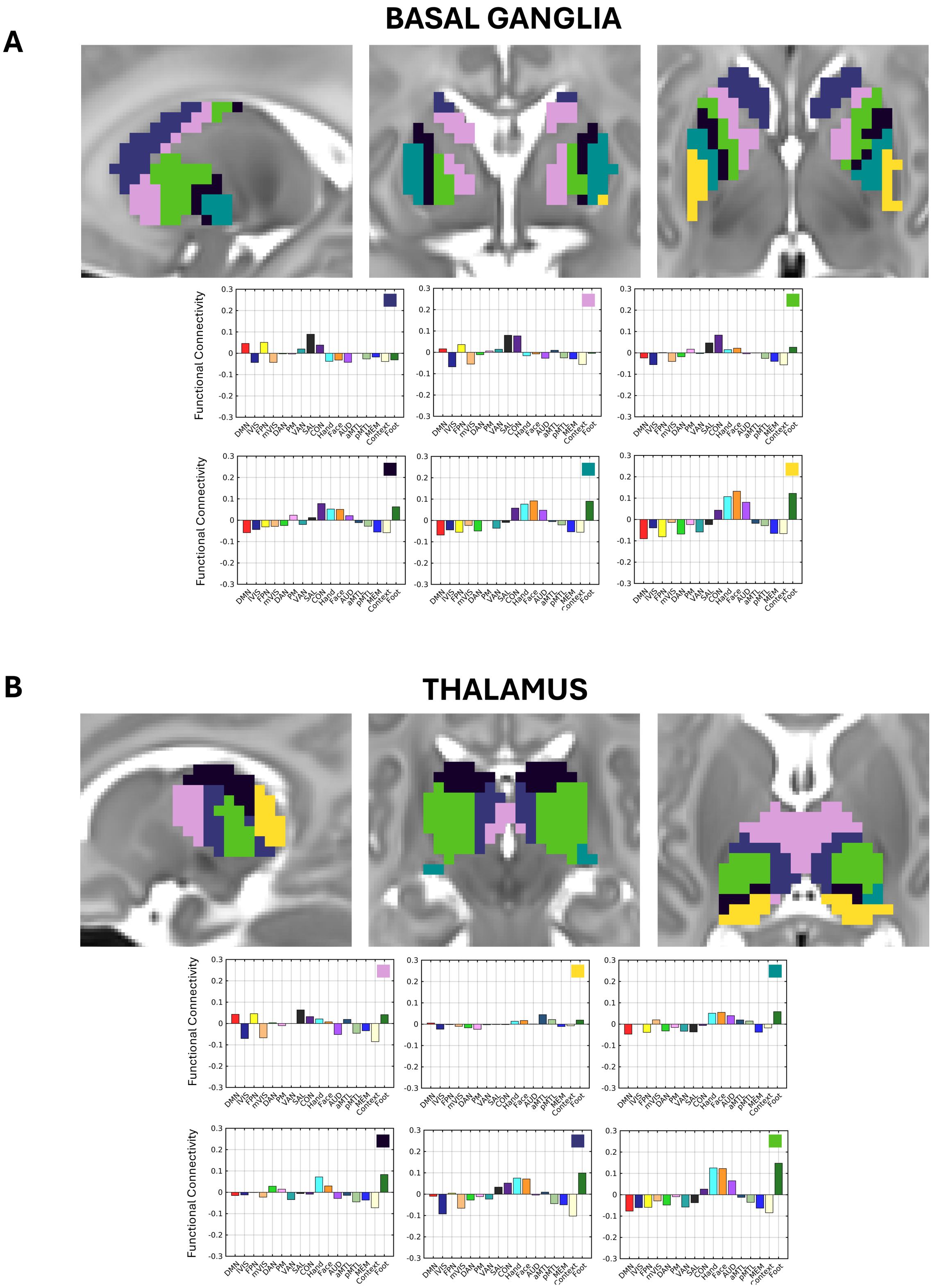


**Supplementary Figure 17.** *K=6 clustering results for the neonatal* ***A****) basal ganglia and* ***B)*** *thalamus according to FC profiles with the Midnight Scan Club networks.* Sagittal (left), coronal (middle), and transverse (right) slices are displayed. Basal ganglia coordinates: x = -20, y = -3, z = 3. Thalamus coordinates: x = -12, y = -24, z = 9. Average network profiles showing average FC to each network for the voxels within each cluster are depicted below. Each network profile graph contains a box with its corresponding cluster color in the top right.

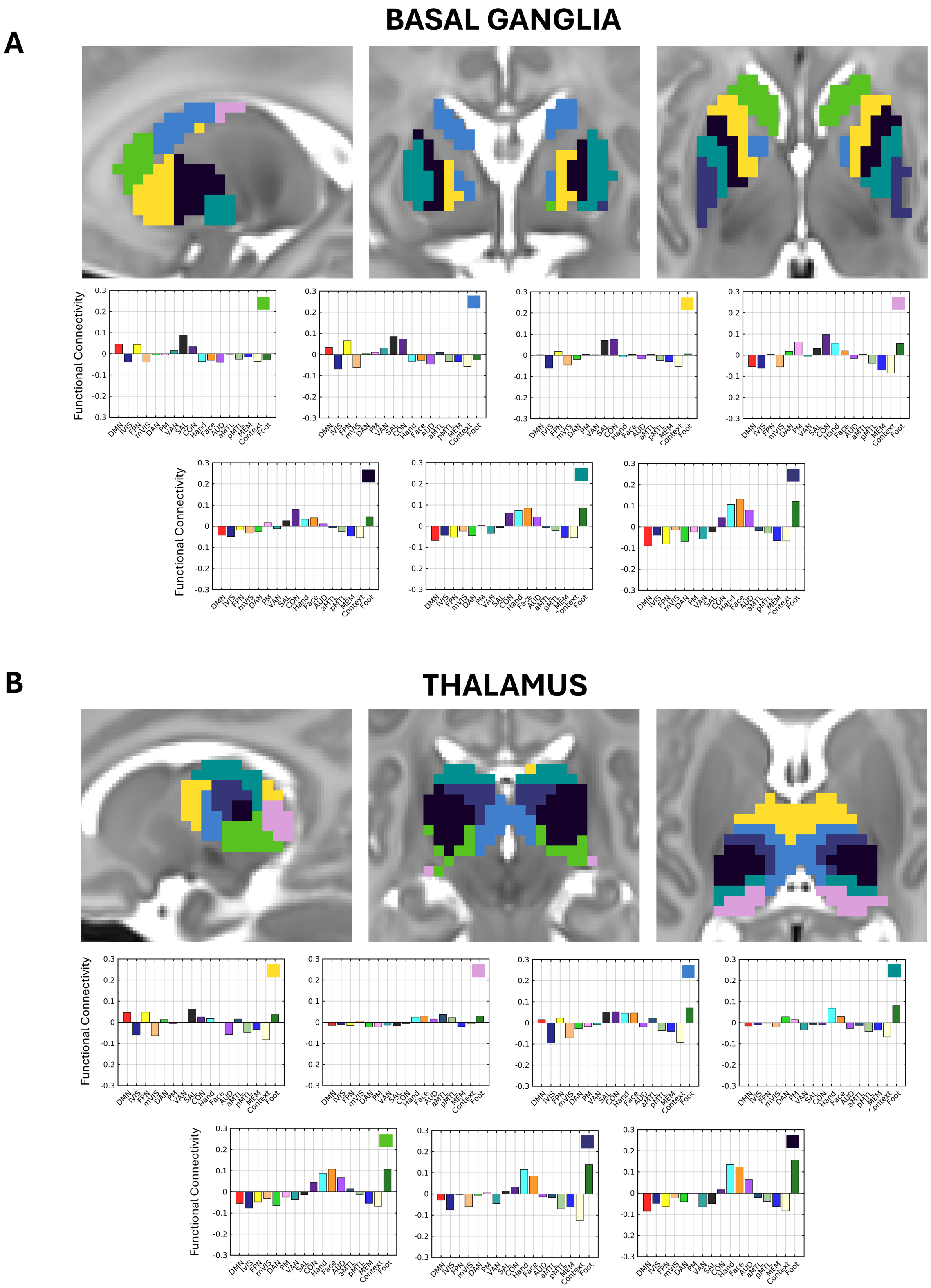


**Supplementary Figure 18***. K=7 clustering results for the neonatal* ***A****) basal ganglia and* ***B)*** *thalamus according to FC profiles with the Midnight Scan Club networks.* Sagittal (left), coronal (middle), and transverse (right) slices are displayed. Basal ganglia coordinates: x = -20, y = -3, z = 3. Thalamus coordinates: x = -12, y = -24, z = 9. Average network profiles showing average FC to each network for the voxels within each cluster are depicted below. Each network profile graph contains a box with its corresponding cluster color in the top right.

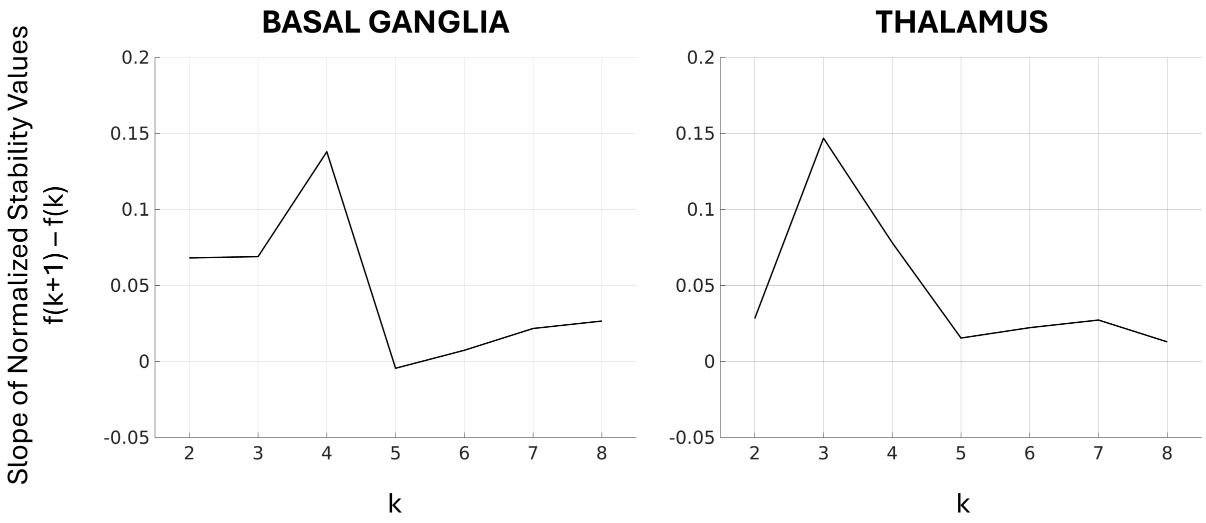


**Supplementary Figure 19***. Cluster stability graphs for k = 2 to k = 9 in the neonatal* ***A) basal ganglia*** *and* ***B)*** *thalamus.* Line plots display the difference between the average normalized stability values for each k, i.e. the slope of stability.

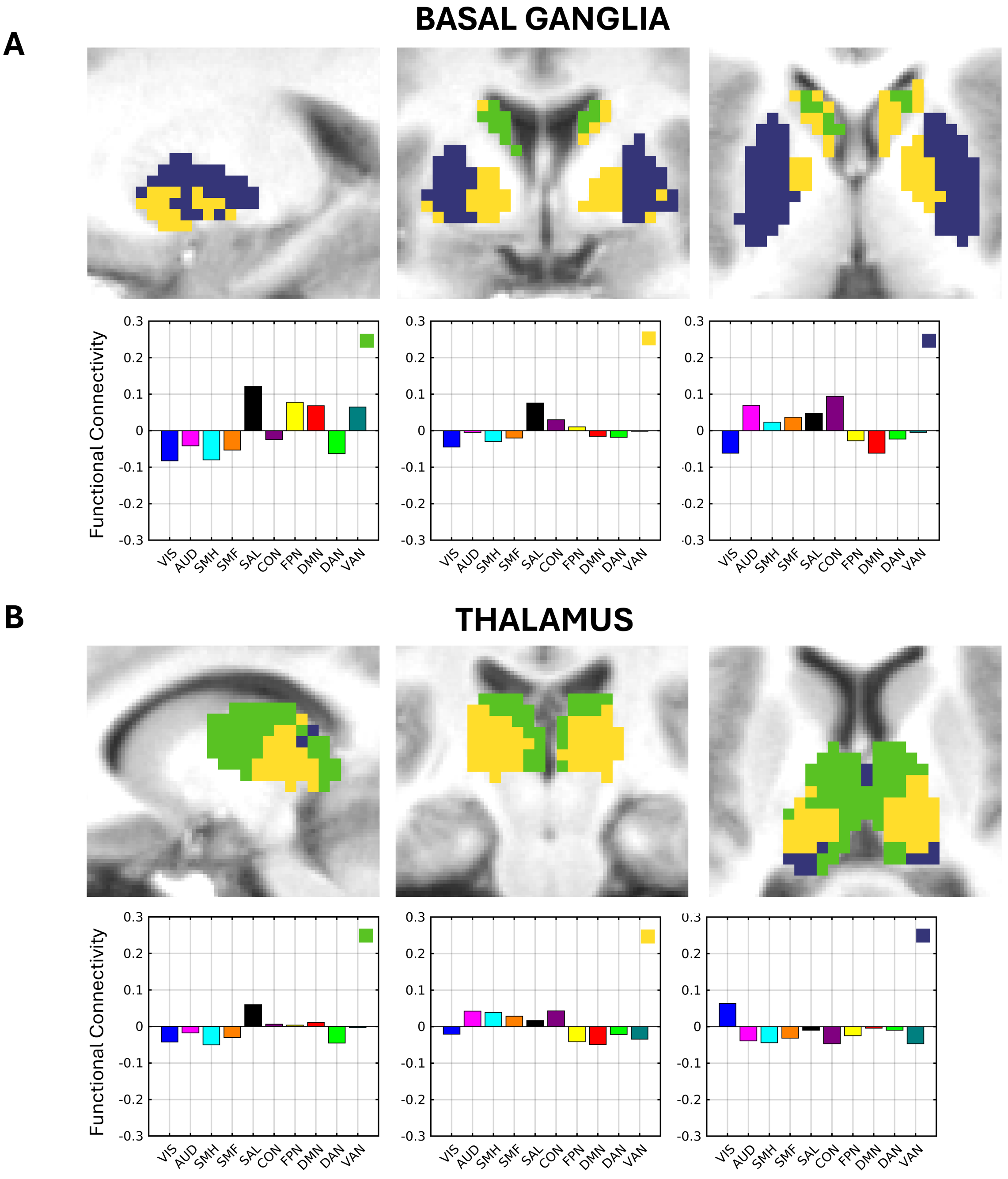


**Supplementary Figure 20.** *K=3 clustering results for the child* ***A****) basal ganglia and* ***B)*** *thalamus according to FC profiles with the Gordon networks.* Sagittal (left), coronal (middle), and transverse (right) slices are displayed. Basal ganglia coordinates: x = -20, y = -3, z = 3. Thalamus coordinates: x = -12, y = -24, z = 9. Average network profiles showing average FC to each network for the voxels within each cluster are depicted below. Each network profile graph contains a box with its corresponding cluster color in the top right.

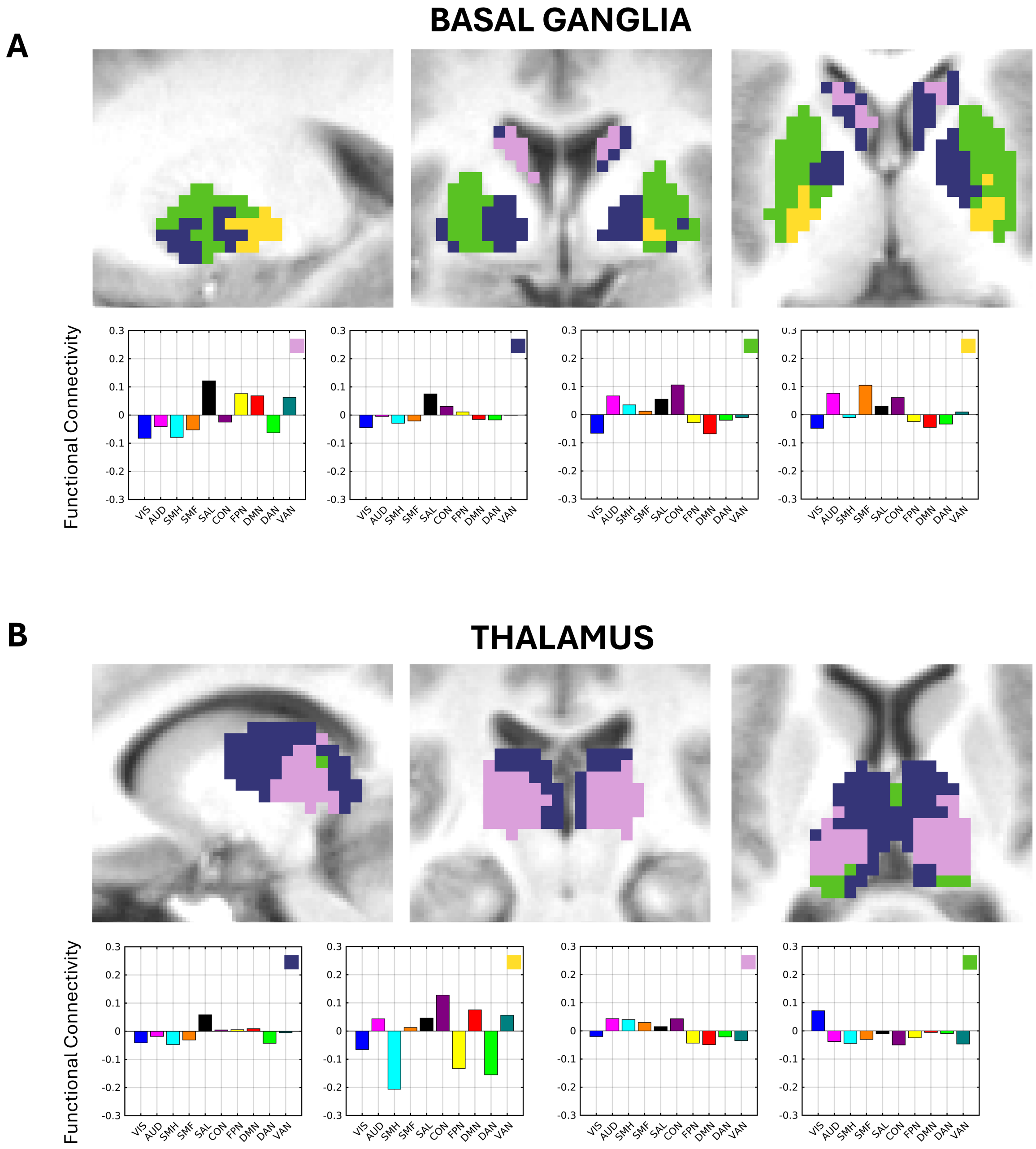


**Supplementary Figure 21***. K=4 clustering results for the child* ***A****) basal ganglia and* ***B)*** *thalamus according to FC profiles with the Gordon networks.* Sagittal (left), coronal (middle), and transverse (right) slices are displayed. Basal ganglia coordinates: x = -20, y = -3, z = 3. Thalamus coordinates: x = -12, y = -24, z = 9. Average network profiles showing average FC to each network for the voxels within each cluster are depicted below. Each network profile graph contains a box with its corresponding cluster color in the top right.

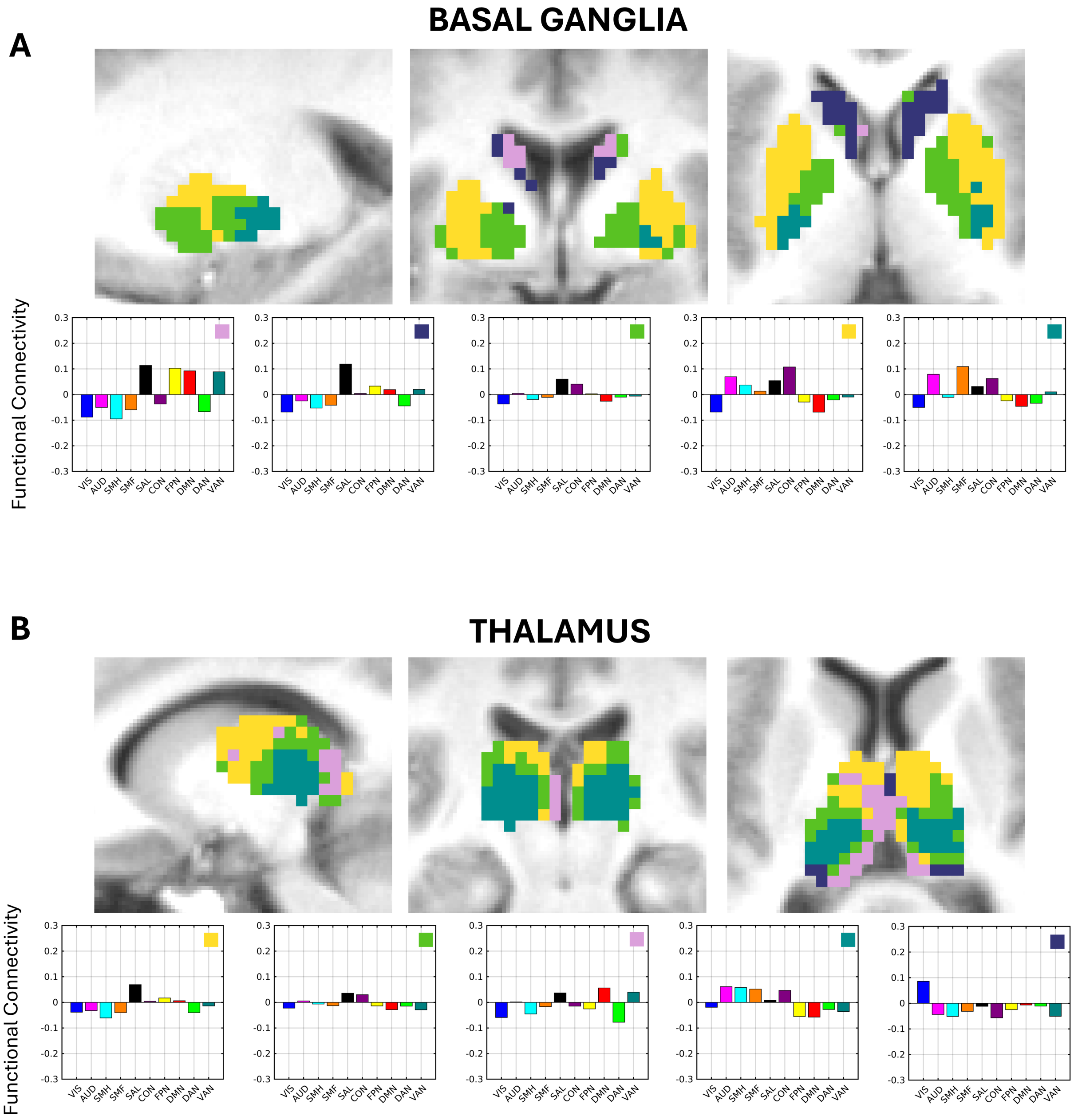


**Supplementary Figure 22.** *K=5 clustering results for the child* ***A)*** *basal ganglia and* ***B)*** *thalamus according to FC profiles with the Gordon networks.* Sagittal (left), coronal (middle), and transverse (right) slices are displayed. Basal ganglia coordinates: x = -20, y = -3, z = 3. Thalamus coordinates: x = -12, y = -24, z = 9. Average network profiles showing average FC to each network for the voxels within each cluster are depicted below. Each network profile graph contains a box with its corresponding cluster color in the top right.

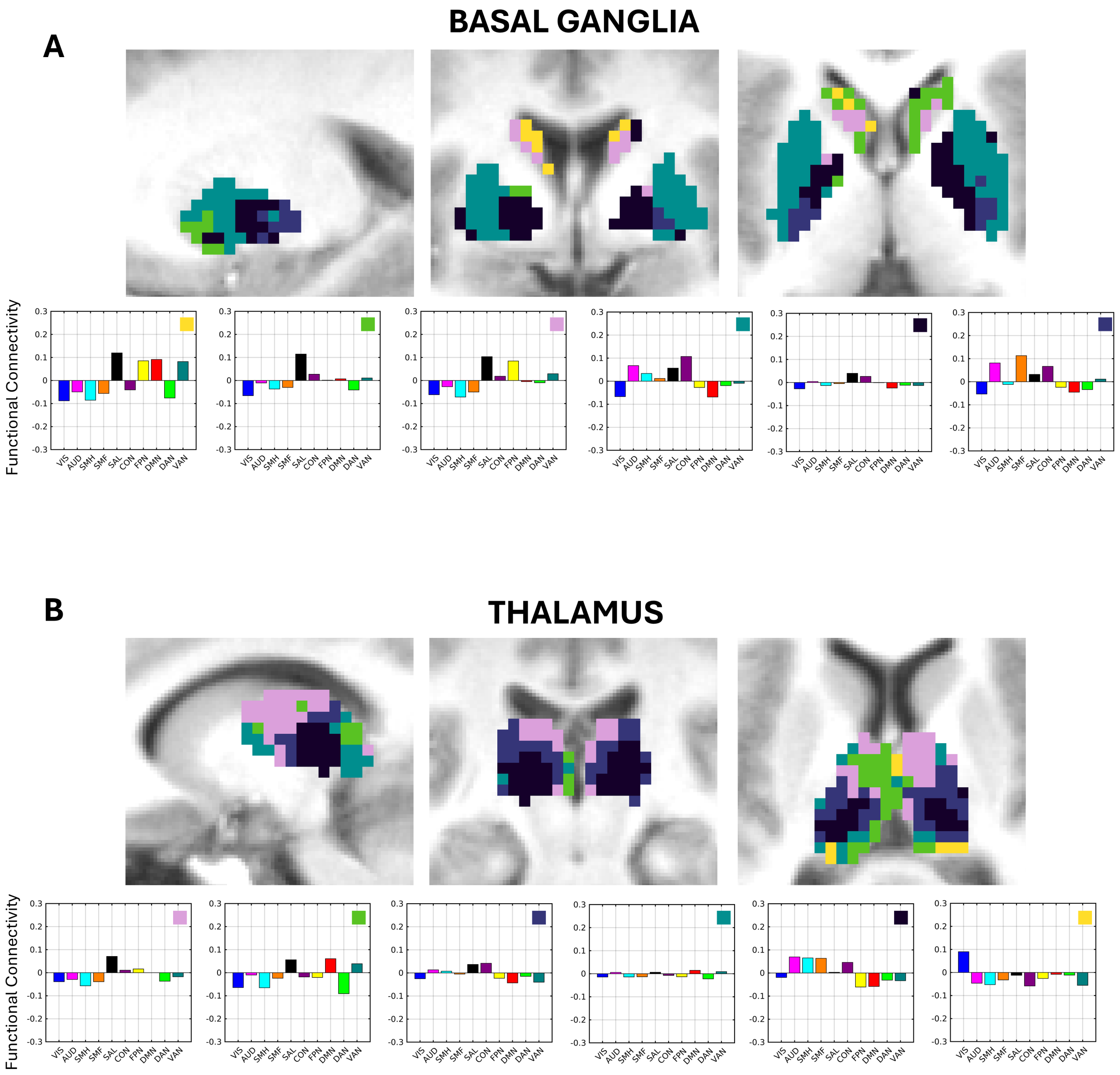


**Supplementary Figure 23***. K=6 clustering results for the child* ***A****) basal ganglia and* ***B)*** *thalamus according to FC profiles with the Gordon networks.* Sagittal (left), coronal (middle), and transverse (right) slices are displayed. Basal ganglia coordinates: x = -20, y = -3, z = 3. Thalamus coordinates: x = -12, y = -24, z = 9. Average network profiles showing average FC to each network for the voxels within each cluster are depicted below. Each network profile graph contains a box with its corresponding cluster color in the top right.

A

**BASAL GANGLIA**

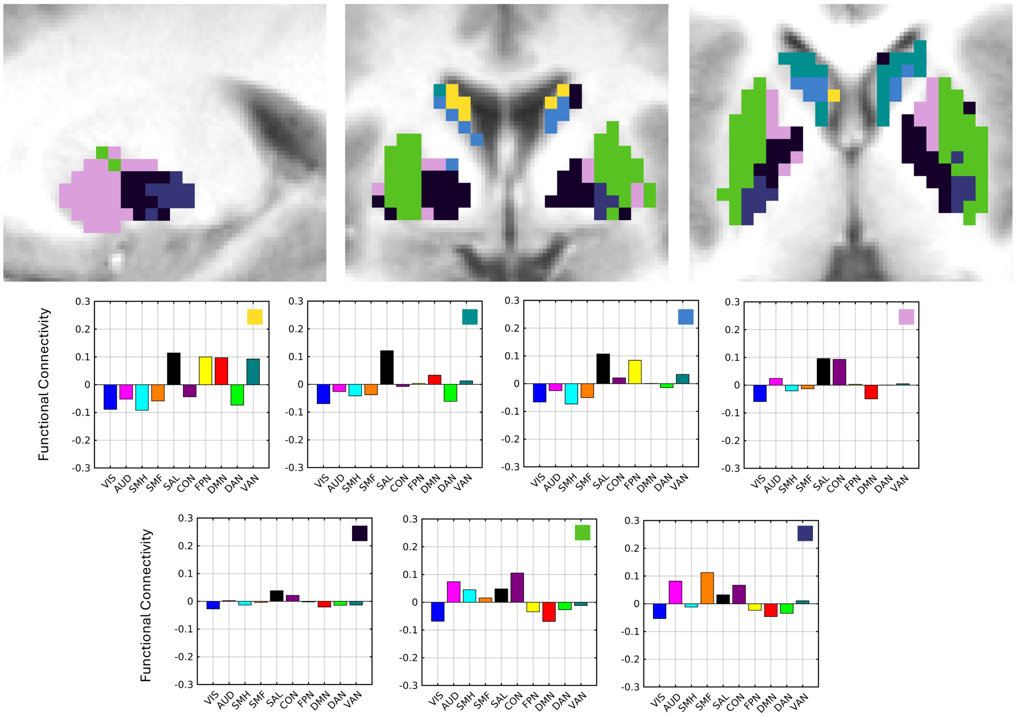


B

**THALAMUS**

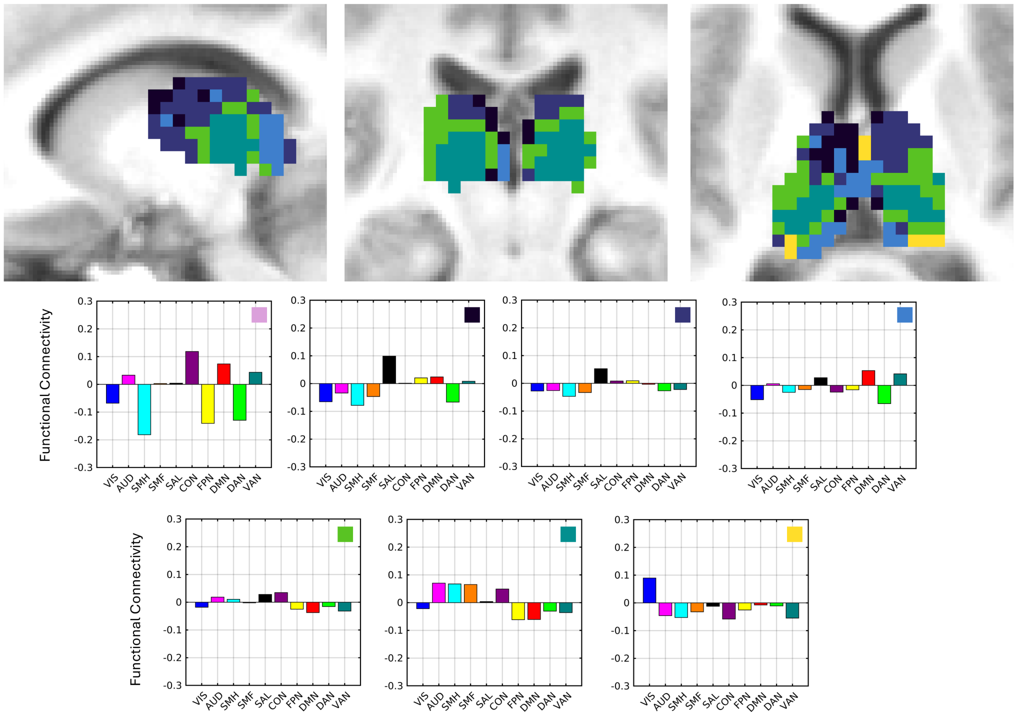

**Supplementary Figure 24*.*** *K=7 clustering results for the child* ***A)*** *basal ganglia and* ***B)*** *thalamus according to FC profiles with the Gordon networks.* Sagittal (left), coronal (middle), and transverse (right) slices are displayed. Basal ganglia coordinates: x = -20, y = -3, z = 3. Thalamus coordinates: x = -12, y = -24, z = 9. Average network profiles showing average FC to each network for the voxels within each cluster are depicted below. Each network profile graph contains a box with its corresponding cluster color in the top right.

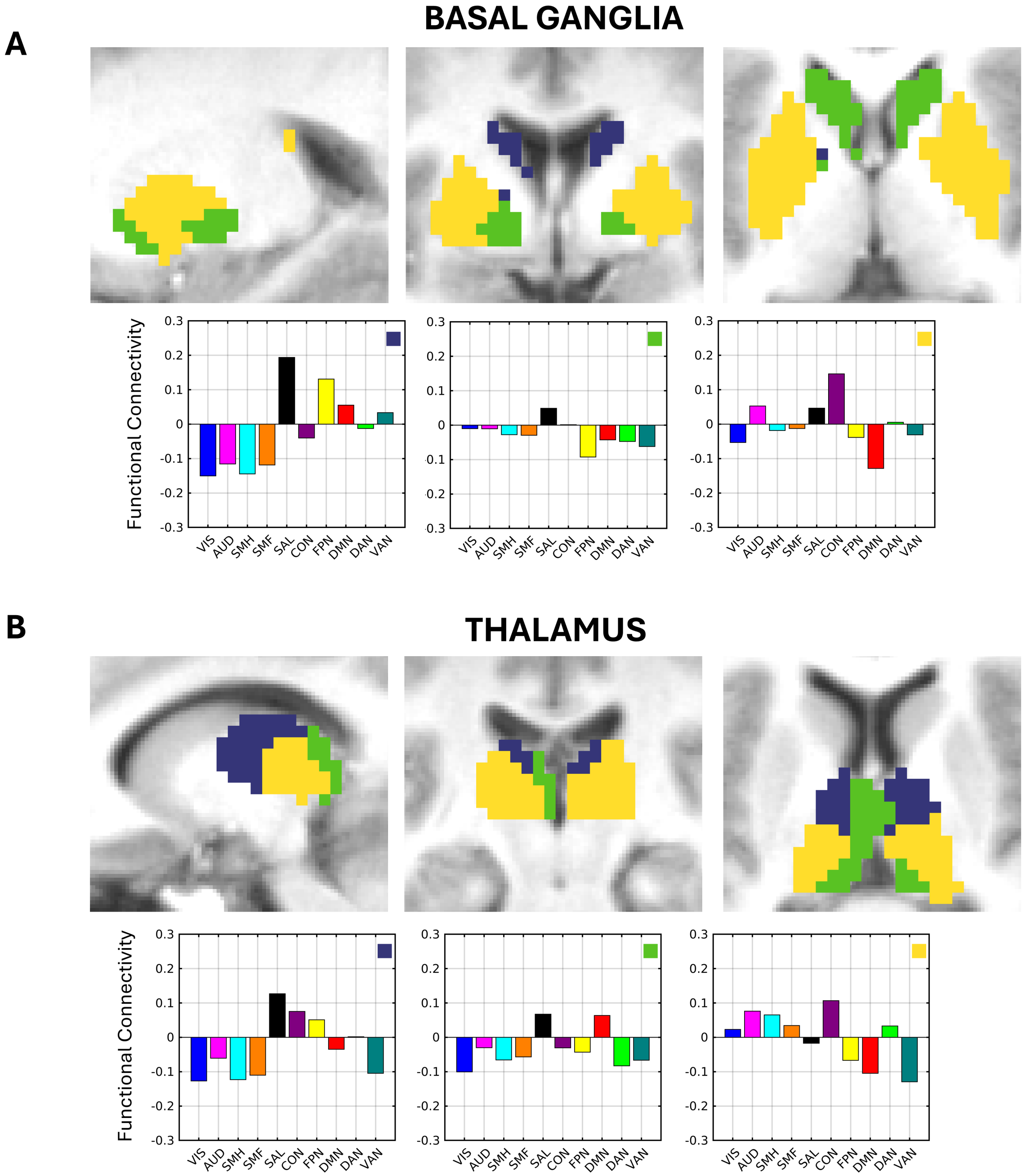


**Supplementary Figure 25.** *K=3 clustering results for the adult* ***A****) basal ganglia and* ***B)*** *thalamus according to FC profiles with the Gordon networks.* Sagittal (left), coronal (middle), and transverse (right) slices are displayed. Basal ganglia coordinates: x = -20, y = -3, z = 3. Thalamus coordinates: x = -12, y = -24, z = 9. Average network profiles showing average FC to each network for the voxels within each cluster are depicted below. Each network profile graph contains a box with its corresponding cluster color in the top right.

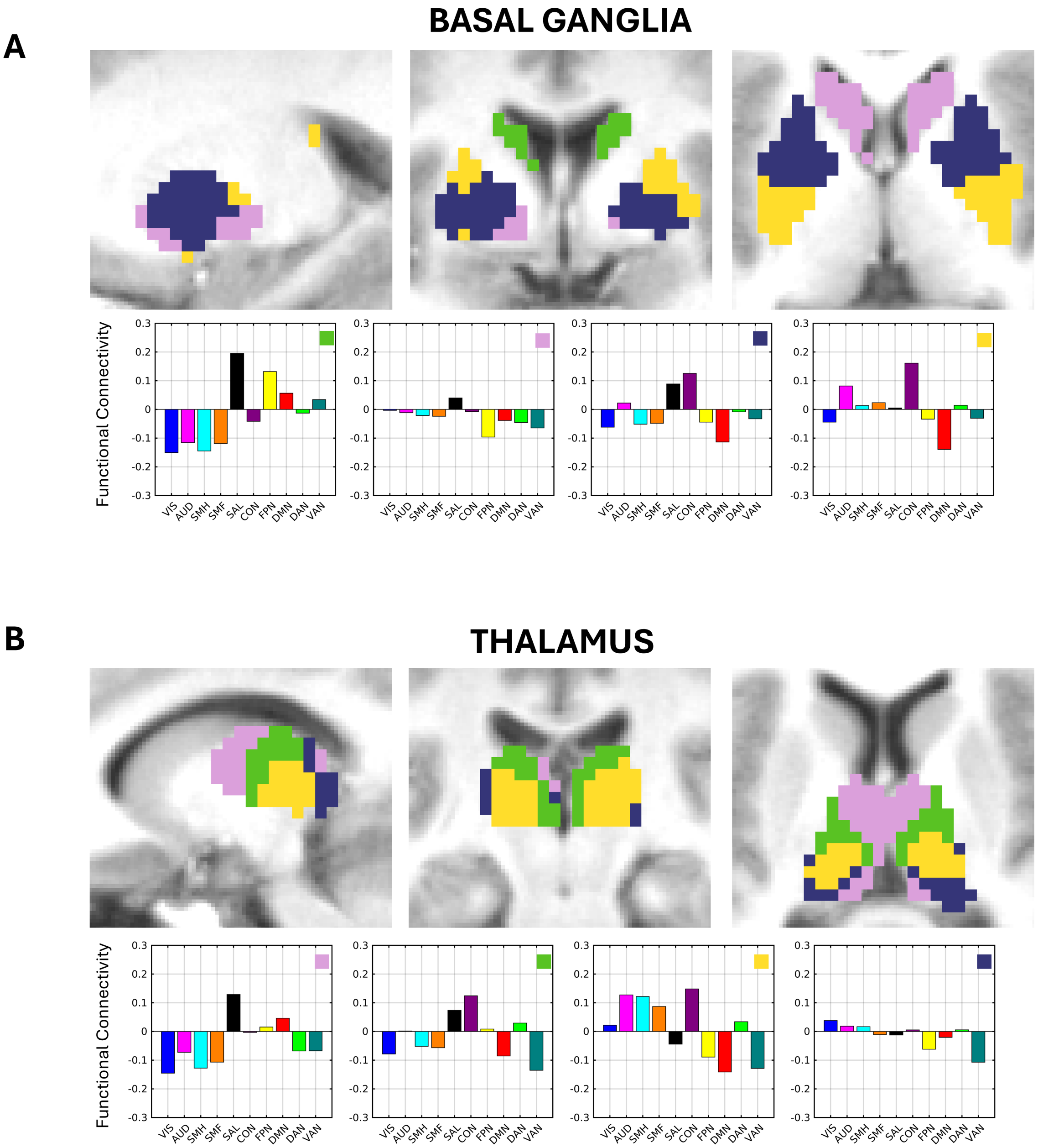


**Supplementary Figure 26.** *K=4 clustering results for the adult* ***A)*** *basal ganglia and* ***B)*** *thalamus according to FC profiles with the Gordon networks.* Sagittal (left), coronal (middle), and transverse (right) slices are displayed. Basal ganglia coordinates: x = -20, y = -3, z = 3. Thalamus coordinates: x = -12, y = -24, z = 9. Average network profiles showing average FC to each network for the voxels within each cluster are depicted below. Each network profile graph contains a box with its corresponding cluster color in the top right.

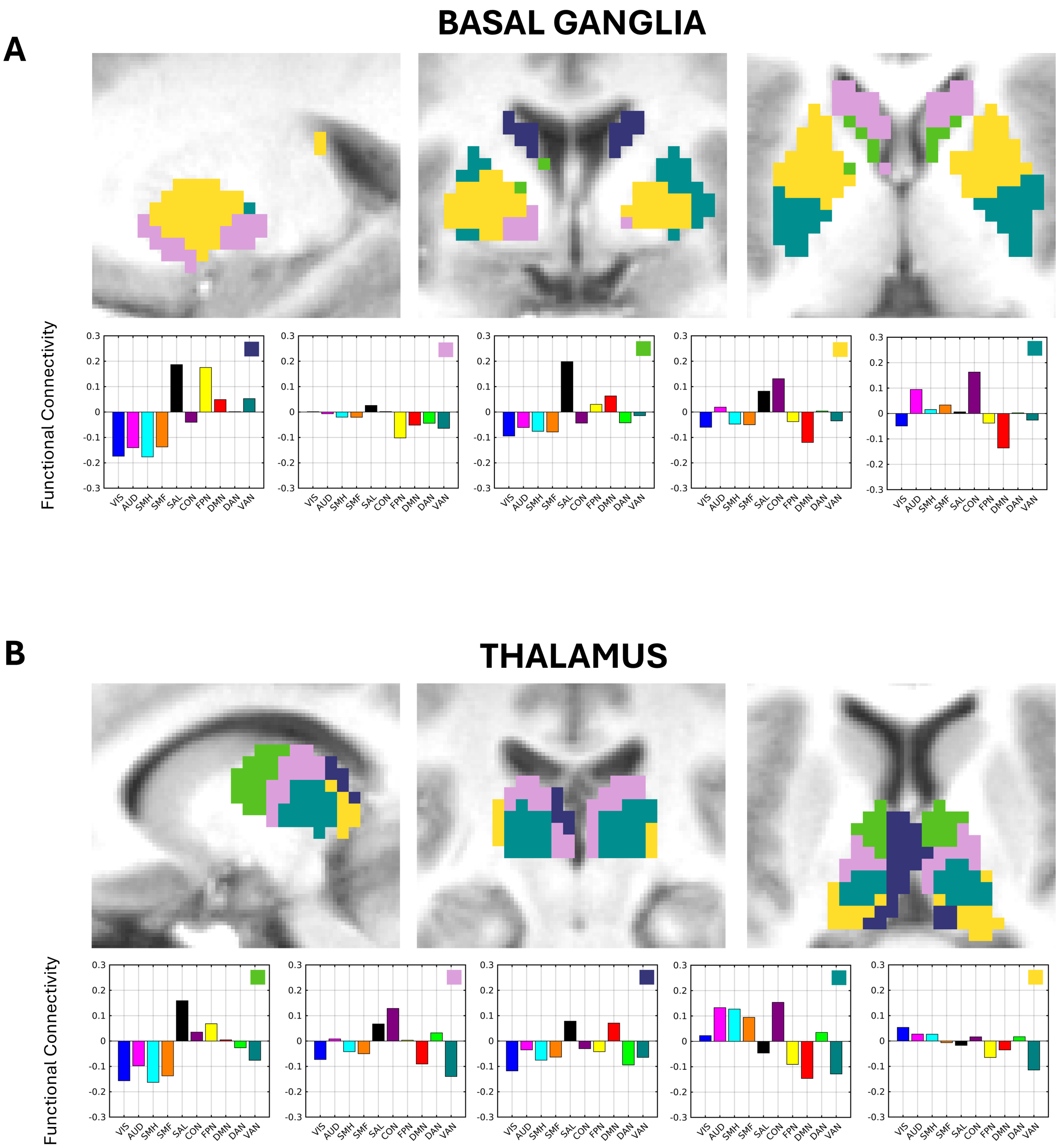


**Supplementary Figure 27*****.*** *K=5 clustering results for the adult* ***A****) basal ganglia and* ***B)*** *thalamus according to FC profiles with the Gordon networks.* Sagittal (left), coronal (middle), and transverse (right) slices are displayed. Basal ganglia coordinates: x = -20, y = -3, z = 3. Thalamus coordinates: x = -12, y = -24, z = 9. Average network profiles showing average FC to each network for the voxels within each cluster are depicted below. Each network profile graph contains a box with its corresponding cluster color in the top right.

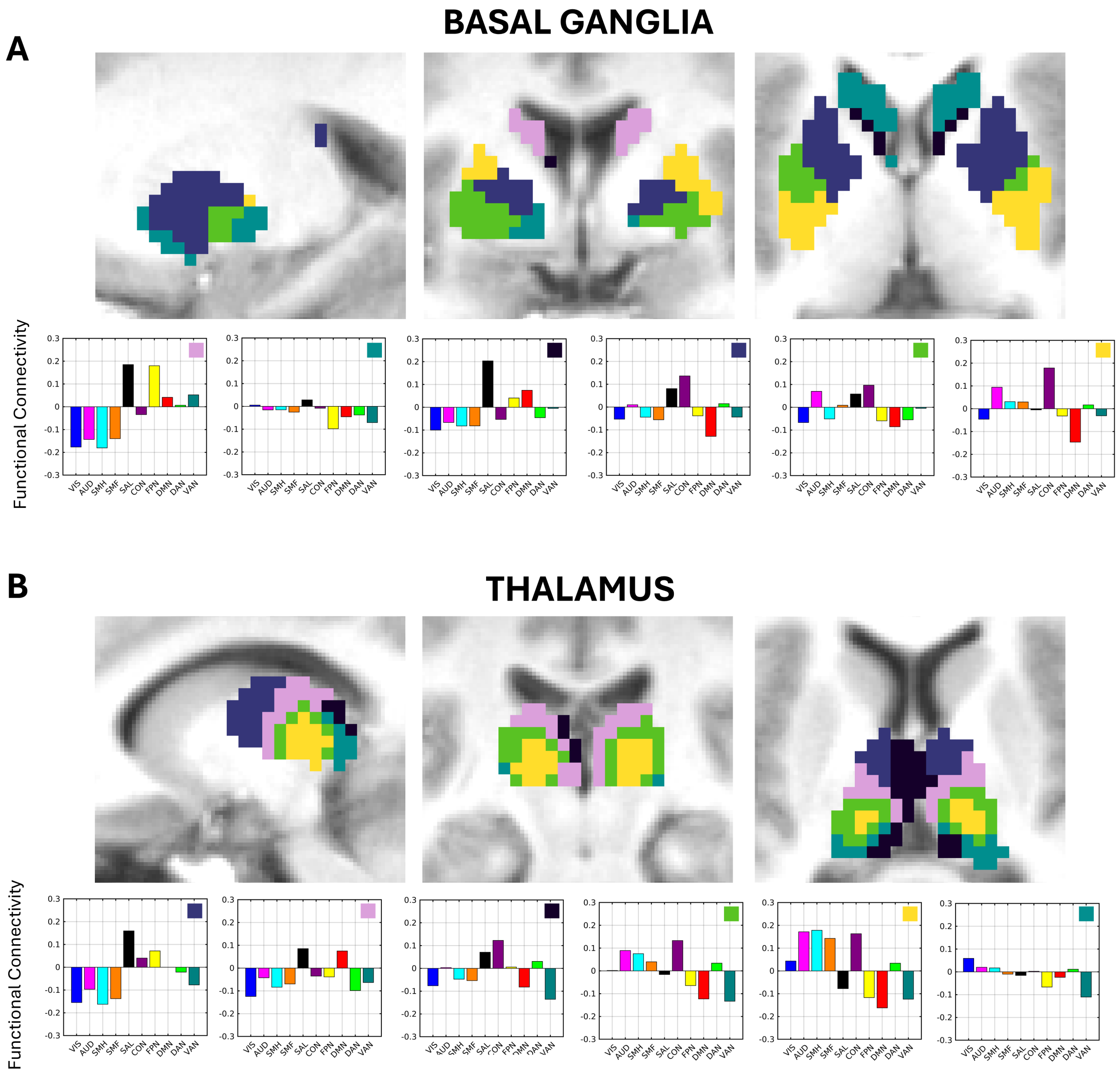


**Supplementary Figure 28***. K=6 clustering results for the adult* ***A****) basal ganglia and* ***B)*** *thalamus according to FC profiles with the Gordon networks.* Sagittal (left), coronal (middle), and transverse (right) slices are displayed. Basal ganglia coordinates: x = -20, y = -3, z = 3. Thalamus coordinates: x = -12, y = -24, z = 9. Average network profiles showing average FC to each network for the voxels within each cluster are depicted below. Each network profile graph contains a box with its corresponding cluster color in the top right.

**A**

### BASAL GANGLIA

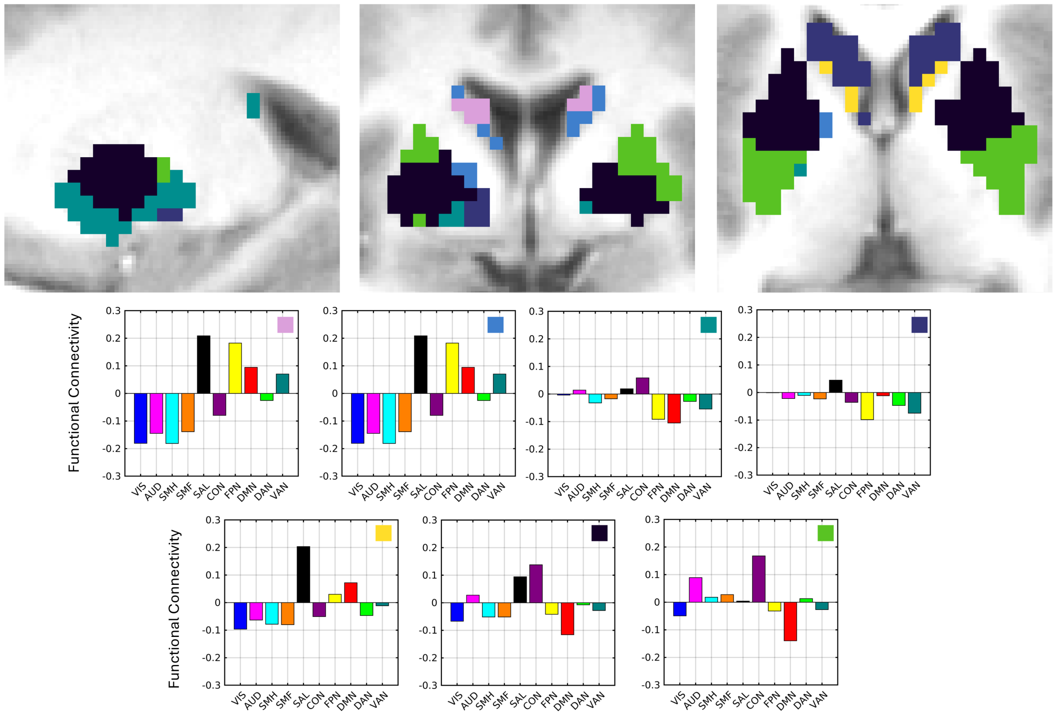


**B**

### THALAMUS

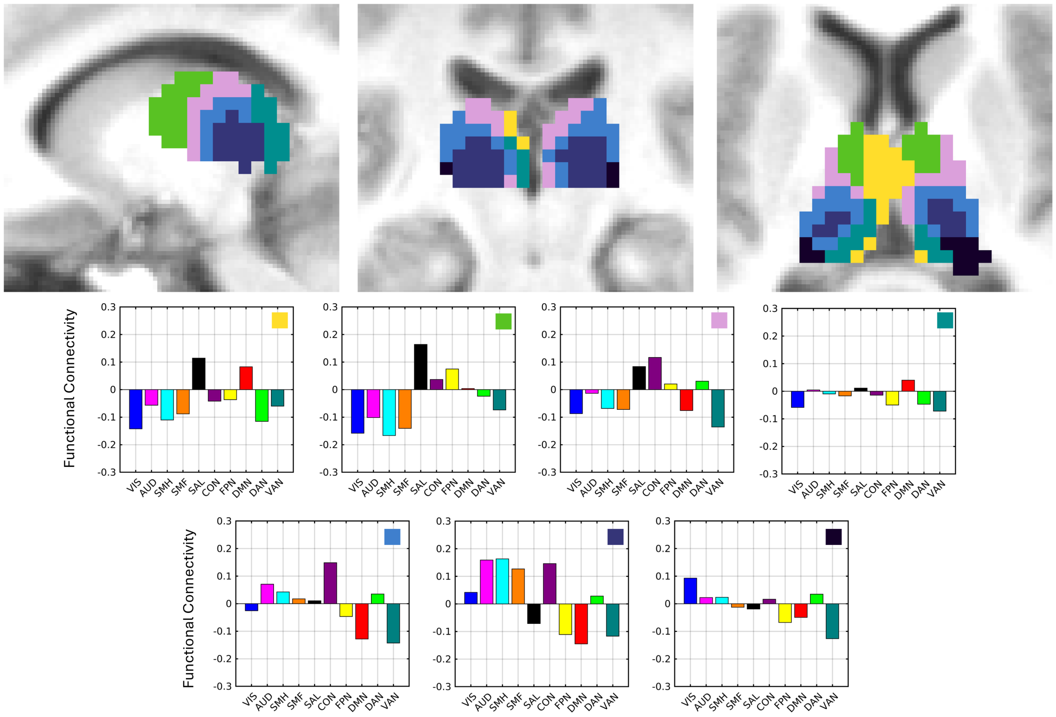

**Supplementary Figure 29***. K=7 clustering results for the adult* ***A****) basal ganglia and* ***B)*** *thalamus according to FC profiles with the Gordon networks.* Sagittal (left), coronal (middle), and transverse (right) slices are displayed. Basal ganglia coordinates: x = -20, y = -3, z = 3. Thalamus coordinates: x = -12, y = -24, z = 9. Average network profiles showing average FC to each network for the voxels within each cluster are depicted below. Each network profile graph contains a box with its corresponding cluster color in the top right.

Marcus, Daniel S., John Harwell, Timothy Olsen, Michael Hodge, Matthew F. Glasser, Fred Prior, Mark Jenkinson, Timothy Laumann, Sandra W. Curtiss, and David C. Van Essen. 2011. “Informatics and Data Mining Tools and Strategies for the Human Connectome Project.” *Frontiers in Neuroinformatics* 5 (June): 4.

Mathur, Amit M., Jeffrey J. Neil, Robert C. McKinstry, and Terrie E. Inder. 2008. “Transport, Monitoring, and Successful Brain MR Imaging in Unsedated Neonates.” *Pediatric Radiology* 38 (3): 260–64.

Myers, Michael J., Alyssa K. Labonte, Evan M. Gordon, Timothy O. Laumann, Jiaxin C. Tu, Muriah D. Wheelock, Ashley N. Nielsen, et al. 2024. “Functional Parcellation of the Neonatal Cortical Surface.” *Cerebral Cortex (New York, N.Y.: 1991)* 34 (2). https://doi.org/10.1093/cercor/bhae047.

Power, Jonathan D., Kelly A. Barnes, Abraham Z. Snyder, Bradley L. Schlaggar, and Steven E. Petersen. 2012. “Spurious but Systematic Correlations in Functional Connectivity MRI Networks Arise from Subject Motion.” *NeuroImage* 59 (3): 2142–54.

Power, Jonathan D., Kelly Anne Barnes, Abraham Z. Snyder, Bradley L. Schlaggar, and Steven E. Petersen. 2013. “Steps toward Optimizing Motion Artifact Removal in Functional Connectivity MRI; a Reply to Carp.” *NeuroImage* 76 (August): 439–41.

Power, Jonathan D., Anish Mitra, Timothy O. Laumann, Abraham Z. Snyder, Bradley L. Schlaggar, and Steven E. Petersen. 2014. “Methods to Detect, Characterize, and Remove Motion Artifact in Resting State fMRI.” *NeuroImage* 84 (January): 320–41.

Smith, Stephen M., Mark Jenkinson, Mark W. Woolrich, Christian F. Beckmann, Timothy E. J. Behrens, Heidi Johansen-Berg, Peter R. Bannister, et al. 2004. “Advances in Functional and Structural MR Image Analysis and Implementation as FSL.” *NeuroImage* 23 Suppl 1: S208-19.

Sylvester, Chad M., Qiongru Yu, A. Benjamin Srivastava, Scott Marek, Annie Zheng, Dimitrios Alexopoulos, Christopher D. Smyser, et al. 2020. “Individual-Specific Functional Connectivity of the Amygdala: A Substrate for Precision Psychiatry.” *Proceedings of the National Academy of Sciences of the United States of America* 117 (7): 3808–18.

Talairach, Jean, and Pierre Tournoux. 1988. *Co-Planar Stereotaxic Atlas of the Human Brain: 3-Dimensional Proportional System : An Approach to Cerebral Imaging*. New York, NY: Thieme-Stratton.
